# Juvenile social isolation induces gene expression alterations and histone modification dysregulations in nucleus accumbens (NAc) neurons of female mice

**DOI:** 10.64898/2026.05.11.724160

**Authors:** Junpeng You, Akira Uematsu, Asuka Jouji-Nishino, Mai Saeki, Yusuke Kishi

**Affiliations:** Laboratory of Molecular Neurobiology, Institute for Quantitative Biosciences, The University of Tokyo, Tokyo 113-0032, Japan; Laboratory of Molecular Biology, Graduate School of Pharmaceutical Sciences, The University of Tokyo, Tokyo 113-0033, Japan; Human Informatics and Interaction Research Institute, National Institute of Advanced Industrial Science and Technology (AIST), Tsukuba 305-8566, Japan; Department of Animal Resource Sciences, Graduate School of Agricultural and Life Sciences, The University of Tokyo, Tokyo 113-8657, Japan; Division of Aging Biology, Research Institute for Science and Technology, Tokyo University of Science, Chiba 278-8510, Japan

**Keywords:** Juvenile social isolation, Nucleus accumbens (NAc), H3K4me1, H3K4me3, H3K27ac, H3K27me3, Brd4, Setd1a, Kdm6b

## Abstract

Lack of social interaction results in various behavioral abnormalities in rodents, including increased anxiety levels, altered sociability, and impaired cognitive ability. Epigenetic factors regulate gene expression, however, how they are involved in juvenile social isolation (jSI)-induced behavioral alterations remains largely unknown. Here, we focused on the nucleus accumbens (NAc), a critical brain region of the reward system that regulates motivation-related behaviors. We first performed RNA-seq on neuronal nuclei and found alterations in genes related to neuronal function, as well as in transcriptional and epigenetic regulation. Protein-protein interaction (PPI) analysis of differentially expressed genes (DEGs) showed that top key nodes among down-regulated genes include membrane receptors (*Ntrk2*, *Grin3a*, and *Grik1*) and an apoptosis regulator (*Bcl2*). To further investigate the association between jSI-induced gene expression alterations and histone modifications, we next performed CUT&Tag for four histone modifications (H3K4me1, H3K4me3, H3K27ac, and H3K27me3), and the results implied that epigenetic alterations may also play a role in neuronal function as well as transcriptional regulation. Reanalysis of previously published RNA-seq data on the manipulation of histone modification-associated factors (including Kdm6b, Brd4, and Setd1a) suggested that these enzymes are candidate regulators of jSI-induced gene expression alterations. Taken together, our comprehensive analysis characterizes the association between histone modification and jSI-related alterations of gene expression in the NAc of female mice.

## Introduction

Social isolation (SI) is a form of environmental stress for both humans and rodents. For humans, SI is closely related to loneliness: loneliness refers to a subjective feeling of social disconnection, while SI describes an objective state of lacking social contact. Previous studies on humans have reported that loneliness is associated with various diseases, including depression, post-traumatic stress disorder, and schizophrenia (Liang et al., 2024; Matthews et al., 2019; Taylor, 2020), and the relevance among teenagers is particularly worthy of attention (Diehl et al., 2018; Shah & Househ, 2023). Studies conducted during the COVID-19 pandemic, when social contact was widely restricted, provided further evidence for negative effects (Koszalinski & Olmos, 2022; Koyama et al., 2021). In addition, social isolation is associated with adverse outcomes, including risk of suicide (Calati et al., 2019; Motillon-Toudic et al., 2022; Zhu et al., 2022). In the case of rodents, socially isolated animal models exhibit a range of behavioral abnormalities, which model aspects of various human diseases, including depression and schizophrenia (Fujiwara et al., 2017; Jiang et al., 2013; Matsumoto et al., 2019).

Adolescence is a critical period of neuronal development that is sensitive to environmental stresses, including juvenile social isolation (jSI) (Makinodan et al., 2012; Yamaguchi et al., 2024). The effects of jSI on motor, emotional, learning, and sociability-related behaviors in rodents have been widely reported, including hyperactivity, elevated anxiety, impaired spatial learning, and altered play (Li et al., 2021; Powell & Swerdlow, 2023; Varlinskaya et al., 1999; Walker et al., 2019). Importantly, whereas the behavioral effects of isolation imposed in adulthood are relatively transient, isolation during adolescence produces alterations that persist into adult life even after social contact is restored (Kim et al., 2021; Lee et al., 2026; Musardo et al., 2022; Rivera-Irizarry et al., 2020). We have previously shown that two weeks of isolation from postnatal day 21 (P21) produces a heightened fear response in adult female mice (Sazhina et al., 2025). Importantly, the effects of jSI often differ between sexes: socially isolated adolescent females show increased sociability (Jeon et al., 2023), whereas males tend to become more aggressive (Wang et al., 2022). The effects of jSI therefore need to be examined in a sex-specific manner. These findings indicate that rodent models of jSI provide a useful tool for understanding the pathogenesis and molecular mechanisms of human mental disorders.

The nucleus accumbens (NAc) is a critical component of the brain reward circuitry, and dysfunction of the NAc has been reported in animal models of drug addiction (Zinsmaier et al., 2022). NAc dysfunction is also associated with impaired social interaction (Pomrenze et al., 2022; Shan et al., 2022) and abnormal emotion expression (Gebara et al., 2021). An fMRI study reported hyperactivity of the NAc in individuals experiencing loss (O’Connor et al., 2008). Interestingly, gene expression pattern in the NAc also responds to loneliness in the case of human beings (Santiago et al., 2023): the NAc from lonely individuals showed key differentially expressed genes (DEGs) that are associated with both neurodegenerative and neuropsychiatric diseases, including Alzheimer’s disease (AD), Parkinson’s disease (PD), and major depression disorder. Consistently, RNA-seq data from female jSI rats showed that downregulated DEGs in the NAc are associated with AD and PD (Zhao et al., 2022).

Since the behavioral effects of jSI persist long after the isolation period has ended, environmental stress during this window is likely to leave a lasting molecular trace within affected cells, and epigenetic regulation, which can stably maintain altered transcriptional states, has been proposed as a candidate mechanism. Papale et al. found that 5-hydroxymethylcytosine (5hmC) is altered by early life stress in the hypothalamus of mouse, and differentially hydroxymethylated regions (DhMRs) are related to nervous system development and neuronal differentiation (Papale et al., 2017). Besides DNA methylation, histone modifications in the NAc are also affected by environmental stresses, such as early life stress (ELS, including maternal separation) and social defeat stress (SDS) (Kronman et al., 2021; Torres-Berrío et al., 2024). Methylation of histone H3 lysine 79 (H3K79) is affected by ELS, and manipulation of Dot1l, the enzyme catalyzing this modification, is sufficient to recapitulate or rescue ELS-induced changes in gene expression and behavior in the NAc (Kronman et al., 2021). Mono-methylation of H3K27 in the NAc is similarly altered by ELS and confers lasting susceptibility to stress in adulthood (Kronman et al., 2021; Torres-Berrío et al., 2024).

Among histone modifications, H3K4me1, H3K4me3, H3K27ac, and H3K27me3 are the most widely used marks for annotating enhancers, active promoters, and repressed regions (Wiles & Selker, 2017). Some of these have been examined in the NAc: H3K27me3 manipulation in the NAc alters *Cartpt* expression and sensitivity to rewarding effects of cocaine (Winter et al., 2025), while methylation of H3K4 is responsive to early life stress and cocaine exposure (Feng et al., 2014; Reshetnikov et al., 2021), with manipulation of H3K4me1 in the juvenile NAc altering sensitivity to subsequent stress (Rashford et al., 2026). In contrast, little is known about the roles of H3K4me3 and H3K27ac in the NAc under stress, and how any of these modifications are altered after jSI, or how such alterations relate to gene expression, remains largely unknown.

How are histone modifications in the NAc altered by jSI, and how do these alterations relate to gene expression? In this study, we aimed to reveal the association between neuronal gene expression and histone modifications induced by jSI in the NAc in female mice. Unlike previous bulk tissue analyses, we specifically isolated neuronal nuclei from the NAc to dissect neuron-specific epigenetic landscapes without glial contamination, and then conducted RNA-seq and CUT&Tag of H3K4me1, H3K4me3, H3K27ac, and H3K27me3. For the association between histone modification and transcription alterations under jSI stress, we found that synapse-associated genes were downregulated, and histone modifications were altered around genes associated with neuronal development and gene expression regulation. To reveal the upstream epigenetic factors that regulate histone modifications, we reanalyzed previously published data, and the results indicated that epigenetic factors such as Kdm6b, Brd4, and Setd1a are candidate regulators of jSI-induced gene expression alterations. These data together characterize the association between histone modifications and jSI-induced gene expression alterations, and identify candidate upstream factors for further investigation.

## Results

### Alteration of gene expression related to neuronal function and transcription regulation in the NAc by juvenile social isolation (jSI) stress

To reveal how jSI stress alters the transcriptome of the NAc, we performed RNA-sequencing (RNA-seq). The NAc of isolated female mice (isolation for 2 weeks from P21 and followed by re-grouping for another 2 weeks) and group-housed mice for 4 weeks from P21 were dissected, and neuronal nuclei labeled with an anti-NeuN antibody were isolated by fluorescence-activated cell sorting (FACS). Then, RNA was extracted from the isolated nuclei and subjected to RNA-seq (Fig. 1A). After mapping reads to the mouse genome (mm10), we determined the read counts mapped on gene models. To determine DEGs, we used the edgeR software (Y. Chen et al., 2025). We found that 653 genes were down-regulated, and 598 genes were up-regulated in jSI mice (Fig. 1B, Table S1, *p* < 0.05 and fold change (FC) > 1.2).

**Figure 1.**
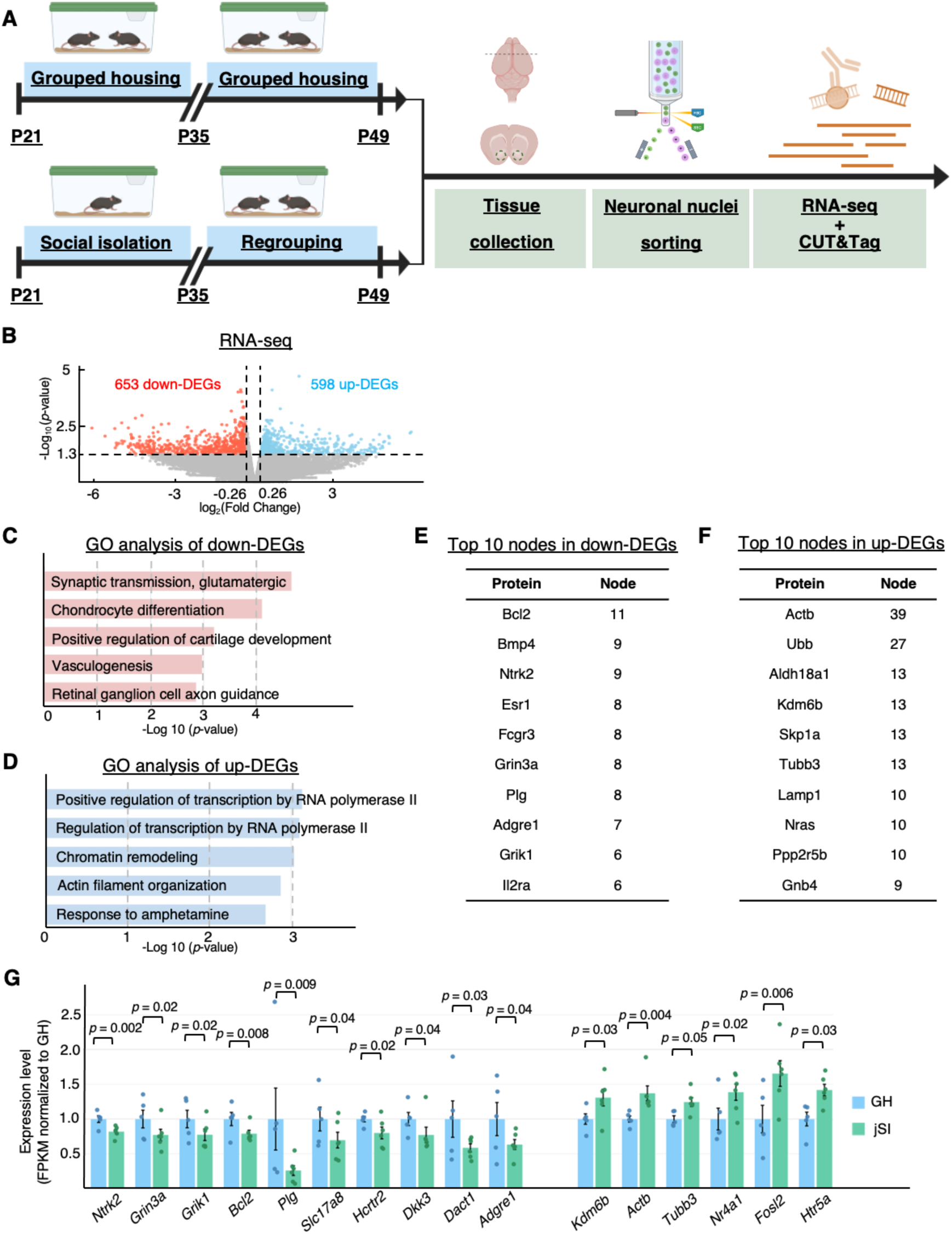
Transcriptome features under juvenile social isolation (jSI) stress. **(A)** Experimental scheme of this study. Mice were either isolated or housed (3-4 mice per cage) in groups from P21 for 2 weeks. After 2 weeks of regrouping, neuronal nuclei, stained with an antibody for NeuN, in the NAc were isolated for both RNA-seq and CUT&Tag. Created in BioRender. Kishi, Y. (2026) https://BioRender.com/j0t4g3g. **(B)** Volcano plot of RNA-seq result. Differentially expressed genes (DEGs) were defined as genes with a *p*-value < 0.05 and Fold Change (FC) > 1.2. Downregulated DEGs (down-DEGs) are labeled as red, and upregulated DEGs (up-DEGs) are labeled as blue. **(C-D)** Gene ontology (GO) analysis of downregulated (C) or upregulated (D) DEGs (down- or up-DEGs). Top 5 GO terms in the “Biological Process” are shown. **(E-F)** Protein-protein interaction (PPI) analysis of down-DEGs (E) or up-DEGs (F). Top 10 nodes are shown. **(G)** Expression levels of the top nodes and other candidate genes in down-DEGs and up-DEGs. GH, grouped housing; jSI, juvenile social isolation; FPKM, fragments per kilobase of transcript per million mapped reads. For RNA-seq, *n* = 6 mice in the jSI group, and *n* = 5 mice in the control group. Data are presented as mean±SEM.

To determine the features of transcriptomic alterations induced by jSI stress, we performed gene ontology (GO) analysis. We found that “synaptic transmission, glutamatergic” is the top GO term of down-regulated DEGs (down-DEGs) (Fig. 1C), and the genes included in this term are: *Grik1*, *Dscam*, *Grin3a*, *Btbd11*, *Slc17a8*, *Grin2d*, and *Ext1*. Up-regulated DEGs (up-DEGs), on the other hand, were enriched with genes related to “regulation of transcription by RNA polymerase II” and “chromatin remodeling” (Fig. 1D). We noticed that several epigenetic factor-encoding genes (such as *Kdm6b*, *Setd1b*, *Setd7*) and transcription factors (such as *Per1* and *Zbtb7a*) are included. These results suggest that jSI stress induced the downregulation of genes related to neuronal function and promoted the expression of genes involved in transcription and epigenetic regulation.

We focused on the potentially important genes within DEGs revealed by protein-protein interaction (PPI) analysis through STRING (Szklarczyk et al., 2023) (https://string-db.org), showing possible interactions among proteins encoded by down-DEGs (Fig. S1) and up-DEGs (Fig. S2). Top nodes in down-DEGs included apoptosis regulator Bcl2 and receptors involved in neuronal functions, such as Ntrk2, Grin3a, and Grik1 (Fig. 1E, G, Fig. S1). Top nodes within up-DEGs included cytoskeletal proteins, such as Actb and Tubb3, and epigenetic factors, such as Kdm6b (Fig. 1F-G, Fig. S2).

These results together suggest that jSI stress affects the expression of both neuronal function and transcriptional regulation-associated genes in the NAc. These genes are downregulated and upregulated, respectively. This indicates their different roles in jSI-mediated behavioral abnormalities.

### Binding of epigenetic regulators around promoters of DEGs in the public database

We next sought to identify candidate upstream transcription factors (TFs) of the DEGs via ChIP-Atlas (Zou et al., 2024) (https://chip-atlas.org), which predicts TFs enriched in the genomic regions of interest based on published ChIP-seq and related datasets. Only data from the “Neural” category under “Cell Class” category were used. Since this database does not include data from the NAc of socially isolated animals, this analysis was intended to generate candidates for further examination rather than to identify regulatory relationships operating in our system.

We analyzed the regions located from 5 kbp upstream to 5 kbp downstream of transcription start sites (TSSs) and found that Foxp1, Cxxc1, Brd4, and Setd1a were enriched at the promoters of down-DEGs (Fig. 2A). The promoters of up-DEGs were predicted to be bound by Ctcf and Foxp1 (Fig. 2B). Among these, Foxp1 is a transcription factor involved in neuronal development (Bacon et al., 2015). Setd1a catalyzes methylation at H3K4 and Cxxc1 is one of its interactors (Tate et al., 2010), while Brd4 recognizes acetylated histones and activates transcription (Jones-Tabah et al., 2022; Mochizuki et al., 2021). Since these factors act through the histone modifications we set out to examine, we focused on the possibility that they contribute to the transcriptional changes observed after jSI.

**Figure 2.**
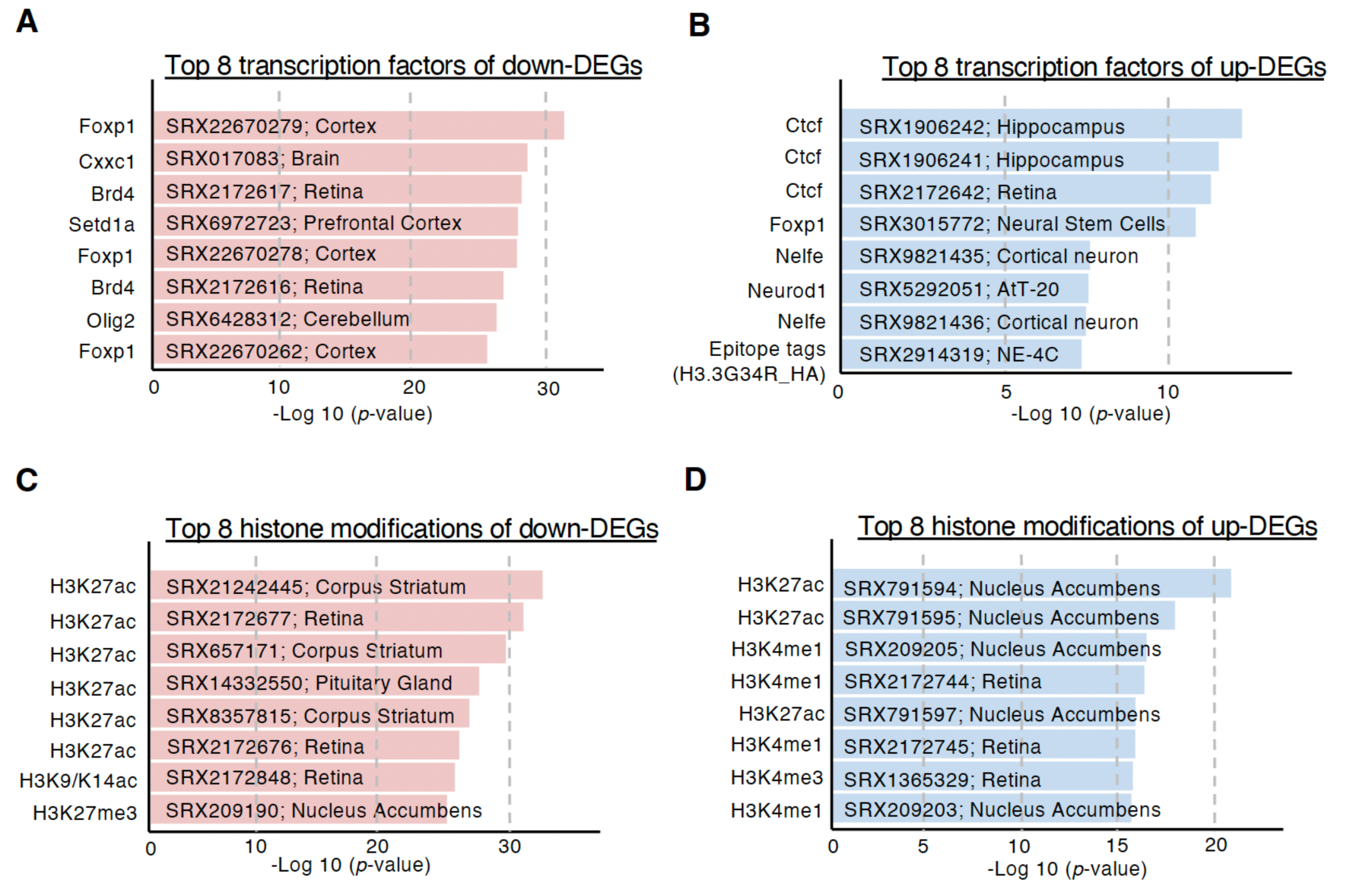
Potential upstream transcription factors of DEGs. ChIP-Atlas analysis for promoter regions (defined as 5 kbp up- and downstream regions of transcription start sites (TSSs)) of DEGs. **(A-B)** Top 8 enriched transcription factors of down-DEGs (A) or up-DEGs (B). The sequencing experiment registered in NCBI SRA database and tissue origin were labeled in the graph. **(C-D)** Top 8 enriched histone modifications of down-DEGs (C) or up-DEGs (D). The sequencing experiment registered in NCBI SRA database and tissue origin were labeled in the graph.

Given that these histone modification-related TFs, such as Kdm6b, Setd1a, and Brd4, were identified as PPI nodes or binding TFs to DEG promoters, we analyzed the histone modifications enriched at promoters of DEGs by ChIP-Atlas. We found that H3K27ac was enriched around the TSS of down-DEGs (Fig. 2C). This is consistent with the TF analysis showing that Brd4 was enriched at these regions. In addition, H3K27ac, together with H3K4me1 and H3K4me3, were enriched around the promoters of up-DEGs (Fig. 2D). These results further imply that histone modifications are associated with alterations of gene expression by jSI stress.

Taken together, this analysis identifies candidate epigenetic regulators that may act upstream of the transcriptional changes we observed after jSI.

### Alteration of histone modifications at DEG loci by jSI stress

Our transcriptome analysis suggested the contribution of epigenetic regulation to alterations of gene expression in the NAc by jSI stress. In addition, previous studies suggested that various histone modifications in the NAc are involved in the response to early life stress and social defeat stress (Kronman et al., 2021; Torres-Berrío et al., 2024). To reveal the changes of the epigenetic state by jSI, we conducted CUT&Tag of isolated neuronal nuclei from the NAc with and without jSI for four histone modifications: three active modifications (H3K4me1, H3K4me3, and H3K27ac) and one repressive modification (H3K27me3). After mapping reads to the mouse genome (mm10), we quantified the signal levels of each histone modification in all 5 kbp bins and determined differentially deposited regions (DDRs) using edgeR (Fig. 3A-D, *p* < 0.05 and FC > 2).

**Figure 3.**
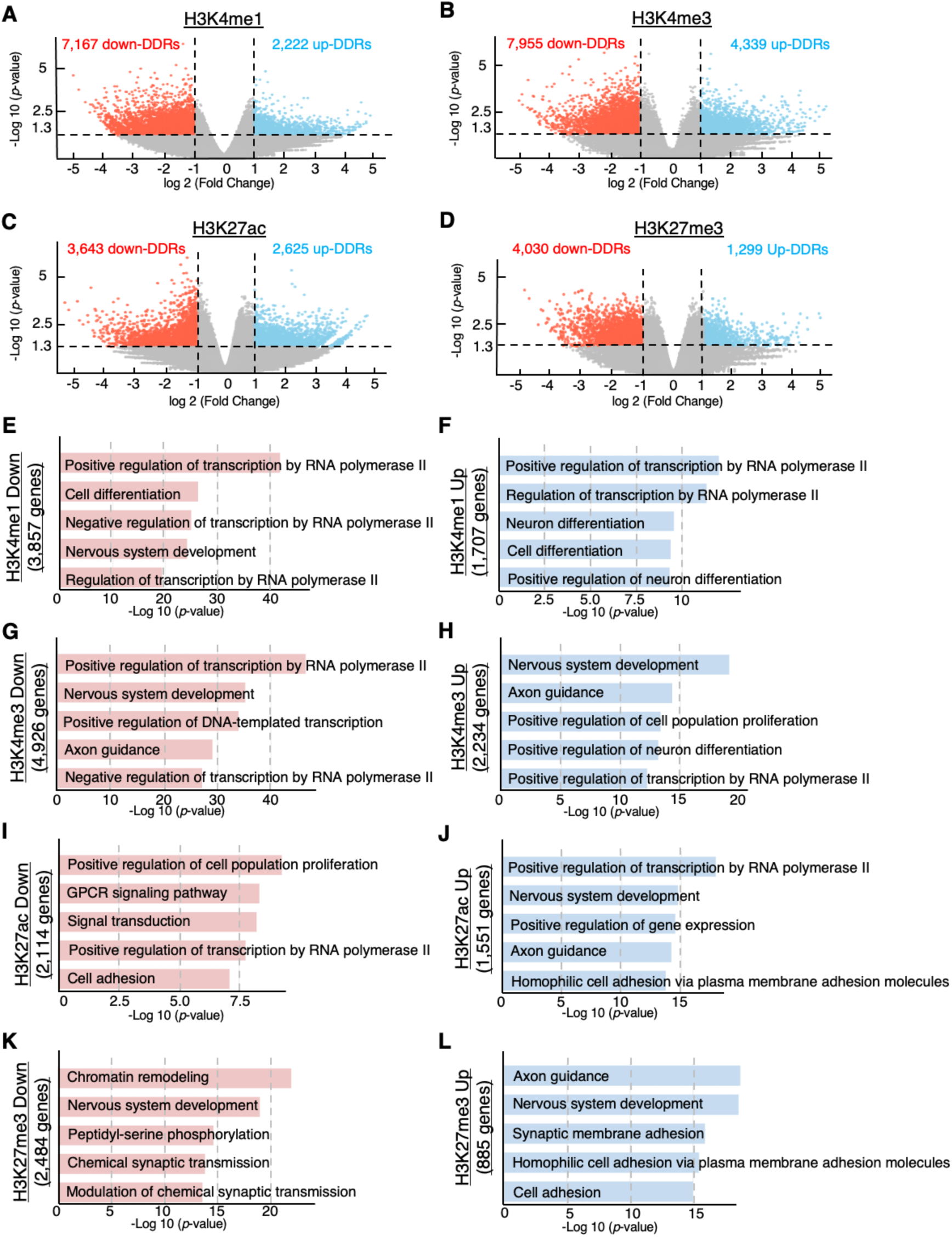
Histone modification distribution alterations under jSI stress. **(A-D)** Volcano plots of the CUT&Tag result for H3K4me1 (A), H3K4me3 (B), H3K27ac (C), and H3K27me3 (D). After quantification of the signals in 5 kbp bins of the whole genome, differentially deposited regions (DDRs) were determined as regions with *p*-value < 0.05 and FC > 2. **(E-F)** Top 5 GO terms (in the “Biological Process”) of down-DDRs (E) and up-DDRs (F) of H3K4me1. **(G-H)** Top 5 GO terms (in the “Biological Process”) of down-DDRs (G) and up-DDRs (H) of H3K4me3. **(I-J)** Top 5 GO terms (in the “Biological Process”) of down-DDRs (I) and up-DDRs (J) of H3K27ac. **(K-L)** Top 5 GO terms (in the “Biological Process”) of down-DDRs (K) and up-DDRs (L) of H3K27me3. For H3K4me1, *n* = 4 mice in jSI, and *n* = 5 mice in control; for H3K4me3, *n* = 6 mice in jSI, and *n* = 8 mice in control; for H3K27ac, *n* = 8 mice in jSI, and *n* = 5 mice in control; for H3K27me3, *n* = 4 mice in jSI, and *n* = 7 mice in control.

To understand the genomic distribution of these DDRs, we next annotated DDRs of each histone modification with their closest genes. GO analysis revealed that two categories of genes were mainly enriched among genes with DDRs across the four histone modifications (Fig. 3E-L). First, genes functionally related to “nervous system development” were enriched among the DDRs of all four histone modifications. The other category included transcription-related terms, such as “regulation of transcription by RNA polymerase II” and “chromatin remodeling”. These results were consistent with the transcriptome analysis that neuronal function and transcription-related genes were affected, and with the transcription factor analysis that epigenetic regulators were predicted to bind to the promoter regions of these genes.

We next examined the association between transcriptomic and epigenetic alterations by jSI. The overlaps between DEGs and DDRs of the four epigenetic modifications were analyzed by Fisher’s exact test (Fig. 4A, Table S1). The results suggest that down-DEGs were associated with changes in H3K4me1, H3K4me3, and H3K27ac in both directions. Consistent with their active roles in transcription, downregulation of H3K4me1 and H3K27ac was more relevant to down-DEGs than up-DEGs, indicating that loss of these active marks accompanies transcriptional downregulation. At the same time, these marks were also increased around a distinct subset of down-DEGs, suggesting that the relationship between histone modification and transcriptional change differs between groups of genes. In contrast, up-DEGs were significantly associated with the downregulation of H3K4me3 and H3K27me3. Since H3K27me3 is a repressive modification, its reduction is consistent with the upregulation of these genes, whereas the concurrent reduction of the active mark H3K4me3 is not, and we therefore consider H3K27me3 the more likely contributor at these loci. These results together indicate that multiple altered histone modifications are associated with global gene expression alterations in NAc neurons, especially for the downregulation of H3K4me1 and H3K27ac.

**Figure 4.**
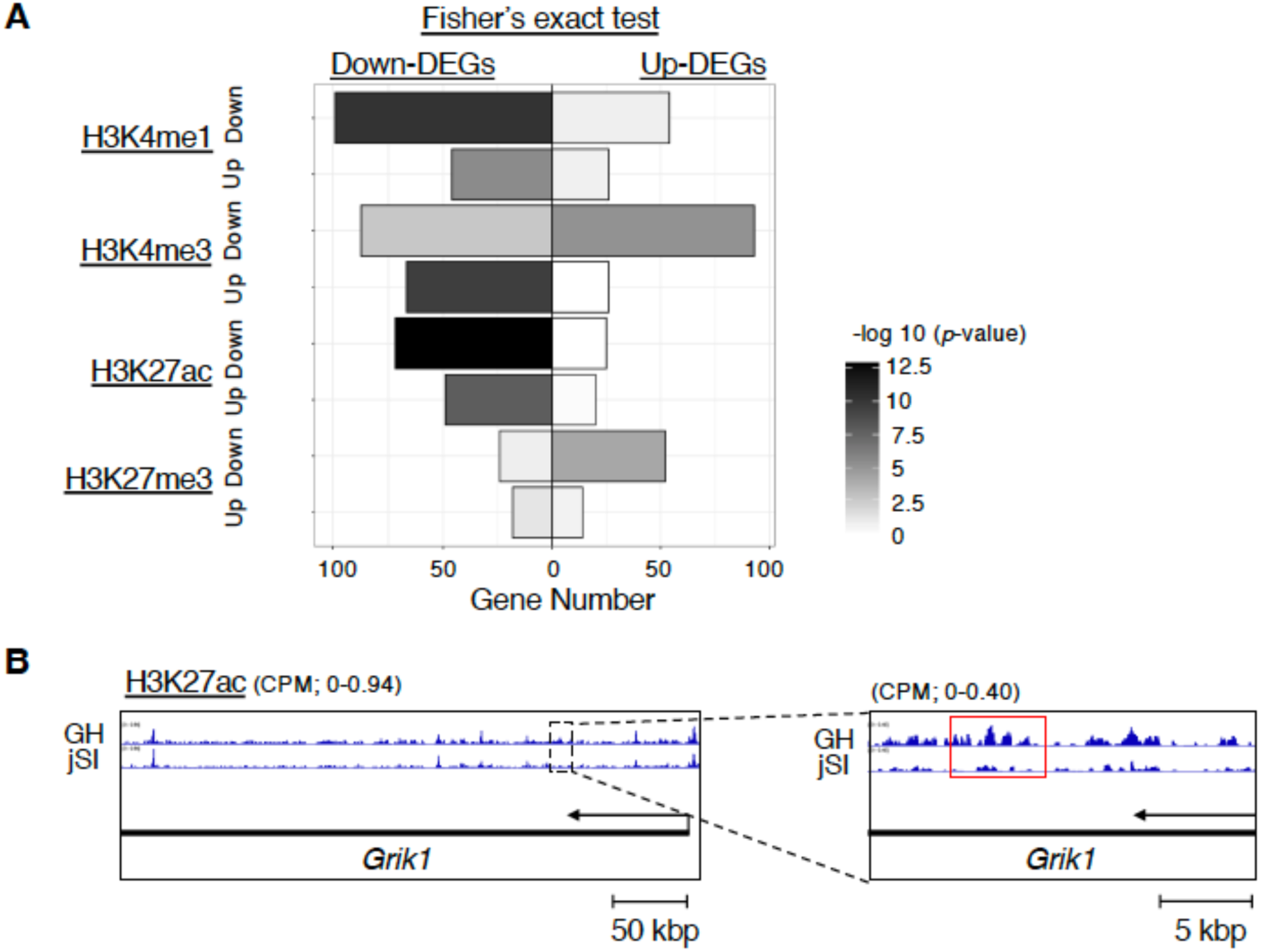
Associations between gene expression and histone modification alterations. **(A)** Overlaps between DEGs and DDRs of four histone modifications. The *p*-values were determined by Fisher’s exact test. **(B)** H3K27ac level around the *Grik1* gene. Averaged counts per million reads (CPM) in each group were visualized by the IGV software (Robinson et al., 2011) (https://igv.org).

To determine whether histone modifications have roles in specific genes, we next focused on the histone modifications at these top nodes revealed by PPI analysis. Among the top 10 nodes of up-DEGs, only *Aldh18a1*, *Lamp1*, and *Gnb4* showed alterations in any of the four histone modifications around their gene loci, all of which were reductions in H3K4me3 (Table S1). By contrast, 8 of the 10 nodes among down-DEGs, except for *Fcgr3* and *Il2ra*, were marked by altered histone modification levels (Table S1). For example, H3K27ac was found to be downregulated around *Grin3a*, *Grik1*, and *Adgre1* (Fig. 4B, Table S1). Among these eight genes, *Bcl2* carried more than one altered modification: downregulated H3K4me1, downregulated H3K4me3, as well as upregulated H3K27me3 were found around *Bcl2* (Table S1). These results imply that the top nodes among down-DEGs are more frequently modulated by multiple histone modifications than those among up-DEGs.

### Possible roles of Kdm6b, Brd4, and Setd1a in the regulation of DEGs in jSI mice

To further investigate the association between histone modifications and gene expression, we analyzed the role of epigenetic regulators on DEGs by reanalyzing published RNA-seq data from neurons. Here, we focused on three histone modification-associated enzymes indicated by our RNA-seq analysis (Kdm6b), and ChIP-Atlas analysis (Brd4 and Setd1a). Among the publicly available data, we selected the datasets in which each enzyme had been perturbed in neural tissue, so that the transcriptional consequences of losing each enzyme could be compared directly with our own DEGs. Since these datasets were obtained under conditions different from ours, we used these comparisons to narrow down candidate genes for further analysis.

Kdm6b demethylates H3K27, and its RNA level was upregulated in jSI mice. Considering the repressive role of H3K27 methylation, upregulation of *Kdm6b* in jSI might contribute to the regulation of up-DEGs. To examine the possible contribution of Kdm6b to jSI, we re-analyzed published RNA-seq data from Kdm6b-knockout mice (Ramesh et al., 2023). The authors performed cerebellum-specific knockout (KO) of Kdm6b by crossing *Atoh1-Cre* and *Kdm6b^fl/fl^* mouse strains and conducted RNA-seq using cerebellar tissue of P14 mice. We re-analyzed their raw data and determined 1302 up-DEGs and 1193 down-DEGs with the same procedure as used for our data (Fig. 5A). We found a significant overlap between up-DEGs by jSI and down-DEGs by Kdm6b KO (*p* = 1.4 × 10^-2^, Fisher’s exact test). Consistently, down-DEGs in jSI were correlated with up-DEGs by Kdm6b KO (*p* = 3.3 × 10^-2^, Fisher’s exact test). Additionally, up- or down-DEGs in jSI were correlated with up- or down-DEGs by Kdm6b KO, respectively (*p* = 8.7 × 10^-4^ between up-DEGs in jSI and up-DEGs in Kdm6b KO, *p* = 1.3 × 10^-4^ between down-DEGs in jSI and down-DEGs in Kdm6b KO, Fisher’s exact test). We next examined the possible downstream genes of Kdm6b in the gene set of up-DEGs in jSI and down-DEGs by Kdm6b KO and found *Nr4a1* and *Fosl2* in this gene set (Fig. 1G, 5G, Table S1). Also, H3K27me3 around the promoter region of *Fosl2* was reduced in jSI (Table S1). *Fosl2* and *Nr4a1* are immediate early genes (IEGs) induced by neuronal activation (Dave et al., 2025; Shi et al., 2024). In addition, *Htr5a*, which encodes serotonin receptor 5A, was also upregulated in jSI and downregulated by Kdm6b KO (Fig. 1G, 5G, Table S1).

**Figure 5.**
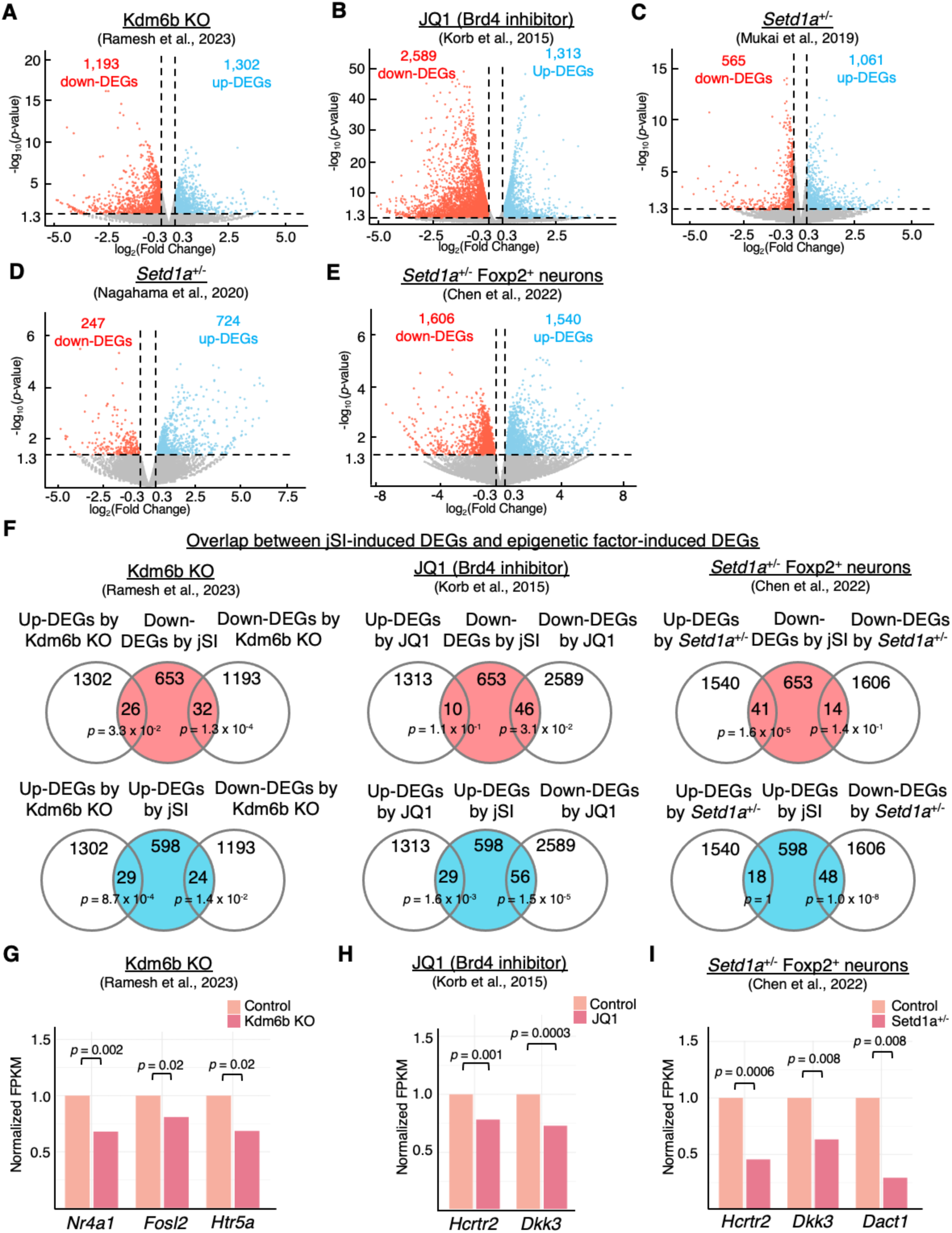
Potential epigenetic factors that regulate DEGs in jSI mice. **(A-E)** Data from the public database were re-analyzed by the same procedure. DEGs were determined as genes with *p*-value < 0.05 and Fold Change (FC) > 1.2. Volcano plots of RNA-seq result published by Ramesh et al. (A), Korb et al. (B), Mukai et al. (C), Nagahama et al. (D), and Chen et al. (E). **(F)** Overlap between jSI-induced DEGs and epigenetic factor-induced DEGs. *p*-values were determined by Fisher’s exact test. **(G-I)** Relative gene expression of potential target genes in Ramesh’s dataset (G), Korb’s dataset (H), and Chen’s dataset (I). Gene expression levels were normalized by the values in the control group. *p*-values were determined by edgeR.

Brd4 was identified as a candidate upstream transcription factor of down-DEGs in jSI, as indicated by ChIP-Atlas analysis (Fig. 2A). Since Brd4 could recognize acetylated histones, including H3K27ac, and activate transcription, we examined whether Brd4 is an epigenetic factor that potentially regulates gene expression in jSI mice. To this end, we analyzed previously published RNA-seq data from primary mouse neurons cultured for 12 days from E16.5 embryonic cortex treated with or without JQ1, an inhibitor for BET family proteins including Brd4, for 24 hours (Korb et al., 2015). We determined 1,313 up-DEGs and 2,589 down-DEGs in JQ1-treated neurons using the same procedure as was used for our data (Fig. 5B). Consistent with the active role of Brd4, down-DEGs by jSI showed significant overlap with down-DEGs by JQ1 treatment (*p* = 3.1 × 10^-2^, Fisher’s exact test), and in addition, up-DEGs by jSI also significantly overlapped with up-DEGs by JQ1 treatment (*p* = 1.6 × 10^-3^, Fisher’s exact test). However, up-DEGs were also significantly overlapped with down-DEGs by JQ1 treatment (*p* = 1.5 × 10^-5^, Fisher’s exact test) (Fig. 5F), indicating that the two gene sets are related but that the direction of change does not correspond in a simple manner. We also examined the possible downstream Brd4 target genes within the gene sets of down-DEGs by jSI and down-DEGs by JQ1 treatment, and we identified *Hcrtr2* and *Dkk3* in these gene sets (Fig. 1G, 5H, Table S1). *Hcrtr2* encodes an orexin receptor, and *Dkk3* inhibits Wnt signaling (Chatterjee et al., 2023; Xiang et al., 2013). These results identify *Hcrtr2* and *Dkk3* as candidate genes whose expression may be regulated by Brd4 in the context of jSI.

Similar to Brd4, Setd1a was also indicated to regulate down-DEGs in jSI. Setd1a is an H3K4 methyltransferase and acts as a transcriptional activator (Kranz & Anastassiadis, 2020). To examine the role of Setd1a, we analyzed three public RNA-seq datasets of Setd1a mutants, which include datasets using the whole prefrontal cortex (PFC) from *Setd1a^+/-^* mice with a *LacZ/Neo* cassette insertion upstream of exon 4 of Setd1a (Mukai et al., 2019); the whole PFC from *Setd1a^+/-^*mice with a frameshift mutation that closely mimics a loss-of-function variant associated with human schizophrenia (Nagahama et al., 2020); and Foxp2-positive neuronal nuclei from the PFC of *Setd1a^+/-^* mice with a frameshift mutation in 15th exon (Chen et al., 2022). We identified 1061 up-DEGs and 565 down-DEGs in Mukai’s data (Fig. 5C), 724 up-DEGs and 247 down-DEGs in Nagahama’s data (Fig. 5D), and 1540 up-DEGs and 1606 down-DEGs in Chen’s data (Fig.5E) using the same procedure as used for our data. All three datasets were compared with our DEGs in the same way. The number of overlapping genes between DEGs in jSI and those in Mukai’s or Nagahama’s data was lower, possibly because of the RNA-seq using the whole PFC that also includes non-neuronal cell types, whereas Chen’s data were obtained from sorted Foxp2-positive neuronal nuclei. In contrast, the overlapped gene set of down-DEGs in jSI and down-DEGs in *Setd1a^+/-^*of Chen’s data included *Hcrtr2* and *Dkk3* as in JQ1-treated primary cortical neurons (Fig. 1G, 5I, Table S1). Consistent with Setd1a’s role in the methylation of H3K4, H3K4me3 levels around *Dkk3* locus were reduced in jSI (Table S1). We also found *Dact1* with H3K4me3 reduction (Table S1). *Dact1* encodes a Wnt signaling-associated scaffold protein (Hou et al., 2015; Jiao et al., 2018). These results identify *Hcrtr2*, *Dkk3*, and *Dact1* as candidate genes whose expression may be associated with Setd1a activity in the context of jSI.

Taken together, these results identify Kdm6b, Brd4, and Setd1a, together with a set of candidate target genes, as starting points for investigating jSI-induced gene expression alterations in the NAc.

## Discussion

### Gene expression dysregulation under social isolation stress

In this study, we analyzed the transcriptome and histone modification distribution patterns in the neuronal nuclei of the NAc from socially isolated mice and group-housed mice. Transcriptome analysis identified 653 downregulated DEGs and 598 upregulated DEGs. Downregulated DEGs were enriched for genes related to neuron-specific functions, such as glutamatergic synaptic transmission. This is consistent with many previous reports that glutamatergic synapses are susceptible to isolation stress (Abramets et al., 2002; Hermes et al., 2011; Hyer et al., 2021; Musazzi et al., 2015). For example, the expression of glutamate receptors, including GluA1 and GluA3, is reduced at protein level in the NAc and caudate putamen (CPu) under chronic social isolation stress (Mao et al., 2022); receptor expression is similarly reduced in the prefrontal cortex after post-weaning isolation (Hermes et al., 2011). Although *Gria1* and *Gria3*, encoding GluA1 and GluA3, respectively, were not among our DEGs, we identified other glutamatergic synapse-associated genes, including *Grin3a* and *Grik1*, suggesting that isolation affects glutamatergic signaling in the NAc through a partly distinct set of genes under our conditions. The NAc receives glutamatergic inputs from the PFC, basolateral amygdala, hippocampus, and ventral tegmental area, and relays this information to the ventral pallidum and other areas of the basal ganglia (Gipson et al., 2014; Goto & Grace, 2008; Walsh et al., 2023). Thus, altered glutamatergic signaling at these synapses may contribute to jSI-induced behavioral abnormalities, such as impaired social interaction, anxiety, and an increased risk of substance abuse. On the other hand, upregulated DEGs were enriched with genes involved in gene expression regulation, including *Kdm6b*, which encodes an epigenetic factor, and these results motivated us to analyze the epigenetic regulators as upstream regulators.

### Histone modification distributions under social isolation stress

TF enrichment analysis revealed that many epigenetic factors (including Foxp1, Brd4, and Setd1a) are enriched around the promoters of down-DEGs. Deficiencies in these factors are associated with distinct behavioral abnormalities. Foxp1 deficiency impairs spatial learning, motor, and communicative behaviors in mice (Araujo et al., 2017; Chasse et al., 2024). Setd1a is a risk gene for schizophrenia in human genetic studies, and its haploinsufficiency recapitulates schizophrenia-related phenotypes in mice (Chen et al., 2022; Mukai et al., 2019; Nagahama et al., 2020). Brd4 has been implicated in cocaine-seeking behavior and in the extinction of fear memory, as well as in Alzheimer’s disease pathology (Guo et al., 2020; Kundakovic, 2022; S. Zhang et al., 2022). Given that Brd4 recognizes acetylated histones such as H3K27ac (Singh & Alauddin, 2023), and Setd1a catalyzes the methylation of H3K4 (Kranz & Anastassiadis, 2020), we hypothesized that histone modifications are involved in jSI-mediated transcriptional alterations. Other evidence from enrichment analysis also indicated that down-DEGs are enriched for specific histone modifications, such as H3K27ac and H3K27me3, while up-DEGs are enriched with H3K27ac, H3K4me1, and H3K4me3. A previous report suggests that the effect of H3K4me1 loss in *SNCA* transgenic mice was partially dampened by enriched environment (Schaffner et al., 2023), and our results indicate that these histone modifications may influence gene expression in the NAc under jSI stress as well.

To verify this hypothesis, we focused on four histone modifications: H3K4me1, H3K4me3, H3K27ac, and H3K27me3. These histone modifications are known as either active markers (H3K4me1, H3K4me3, H3K27ac) or a repressive marker (H3K27me3) of gene expression. Our GO analysis results suggest that all four histone modifications share two common functional groups. The first group is associated with neuronal development, which includes GO terms such as “nervous system development” and “axon guidance”. Since our mice were isolated from P21 to P35, a period critical for the maturation of neurons (Makinodan et al., 2012; Walker et al., 2019; Yamaguchi et al., 2024), it is possible that the neuronal development process is affected by the isolated housing environment. Neurons during adolescence mainly experience synaptic pruning and elimination (Afroz et al., 2016; Germann et al., 2021; Watanabe & Kano, 2024), and isolation stress may impair such processes. Interestingly, development-associated genes may not only function during the developmental stage of life. According to a loneliness study in humans, neuronal development-associated genes are differentially expressed in the NAc (Santiago et al., 2023). In addition, “nervous system development” is the top GO term for both hyper- and hypo-DhMRs in mice that experienced early life stress (Papale et al., 2017). Besides DNA methylation, other epigenetic mechanisms, including histone modifications, are involved in neuronal development as well (Singh et al., 2025). Consistent with these findings, our results indicate that nervous system development-associated genes are susceptible to environmental stress, and that various histone modifications are involved in the stress-responsive regulation of nervous system development.

The second group contains gene expression regulation-associated terms, including “regulation of transcription by RNA polymerase II” and “chromatin remodeling”. Genes associated with transcriptional and chromatin regulation carried altered histone modifications, and since such genes act on the expression of many downstream targets, changes at these loci may have consequences that extend well beyond the genes themselves. This may be particularly relevant during the developmental window examined here, when transcriptional programs are actively shaping neuronal maturation (Jain et al., 2001; Xiang et al., 2020). The abnormal distribution of histone modifications around these regulatory genes may also suggest that the gene expression pattern responding to environmental stimuli is different in isolated animals, which may represent a potential biological mechanism underlying the distinct behavioral patterns in socially isolated mice. We therefore propose that histone modification changes at regulatory genes act as an upstream event after jSI, whose consequences are amplified through the downstream targets of those regulators, a hypothesis that remains to be tested experimentally.

Besides these two main shared functions affected by jSI, our results suggest that other biological processes are potentially mediated by one or more histone modifications. For example, some DDRs of H3K27ac and H3K27me3 are functionally enriched around cell adhesion-associated genes, and this is consistent with previous papers suggesting that cell adhesion is affected by isolation (Santiago et al., 2023; Wu et al., 2022). Many cell adhesion molecules are located in axons and dendrites, and they mediate synapse formation, maintenance, and modulation in normal neurons (Zatkova et al., 2016). Their deficiency causes impaired synapse integrity and abnormal behaviors. Moreover, the previous study on lonely human individuals suggests that the adherens junction pathway is the most affected signaling pathway (Santiago et al., 2023).

To determine whether histone modifications regulate specific genes, we focused on top nodes. *Grik1*, for example, exhibits reduced H3K27ac levels. It encodes a subunit of ionotropic glutamate receptors, and its deficiency has been found in mental diseases, such as schizophrenia and ADHD (Chatterjee et al., 2022; Hirata et al., 2012). The inactivation of *Grik1* in rodents promotes anxiety-like behaviors via glutamatergic transmission (Englund et al., 2021). Our results show that *Grik1* is downregulated after jSI and that H3K27ac around this locus is reduced. We therefore propose reduced H3K27ac contributes to the downregulation of *Grik1*, which in turn may contribute to the anxiety-like phenotypes associated with jSI. Testing this will require manipulating H3K27ac at the *Grik1* locus and assessing the resulting transcriptional and behavioral changes. In addition, several genes are associated with multiple histone modification alterations. *Bcl2*, for example, has downregulated H3K4me1, H3K4me3, and upregulated H3K27me3 levels. *Bcl2* is an apoptosis-related gene that determines neuronal survival under stress. A previous study suggests that chronic social defeat stress decreases the Bcl-2/Bax ratio in NeuN^+^ neurons in the hippocampus (Zhu et al., 2024). Our data suggest that jSI is another type of stress that suppresses *Bcl2* expression, and that the epigenetic factors are candidate upstream regulators. Based on these observations, we propose that the coordinated loss of H3K4me1 and H3K4me3 together with the gain of H3K27me3 around the *Bcl2* locus contributes to its downregulation after jSI, which may in turn affect neuronal survival under stress. The next step is to explore the potential mechanisms of histone modifications regulating glutamatergic synapse-related genes, as well as their roles in controlling behaviors.

### Possible upstream factors of histone modification alteration

To reveal the upstream regulators of histone modifications, we compared our data with previously published data. *Kdm6b* is considered as a risk gene for neurodevelopmental disorders (Rots et al., 2023), and *Kdm6b* activation induces anxiety-like behaviors as well as drug-seeking behaviors (Shentu et al., 2021; Zhang et al., 2018). We found that *Kdm6b* may be at least partially involved in the gene expression changes in jSI mice, especially for candidate genes such as *Nr4a1*, *Fosl2*, and *Htr5a*. *Nr4a1* encodes an immediate early transcription factor that regulates dopamine metabolism, and it has been implicated in drug addiction through changes in histone modification (Carpenter et al., 2020; Engeln et al., 2022; McNulty et al., 2012; Rouillard et al., 2018). This is consistent with reports that jSI induces drug-seeking behaviors (Baarendse et al., 2014; Trezza et al., 2014) and that the NAc is a critical brain region regulating these behaviors (Zinsmaier et al., 2022). *Fosl2*, another immediate early gene, has been reported to be involved in memory maintenance and in Parkinson’s disease (Fan et al., 2020; Mizuno et al., 2020), and *Htr5a*, which encodes serotonin receptor 5A, shows genetic association with schizophrenia (Guan et al., 2016). Based on these observations, we propose that Kdm6b-mediated H3K27 demethylation contributes to the upregulation of *Nr4a1* in the NAc after jSI, a hypothesis that remains to be tested by NAc-specific manipulation of Kdm6b. Future work is necessary to confirm the contribution of Kdm6b protein levels to the candidate genes, including *Nr4a1*.

Brd4 is a potential upstream transcription factor that regulates gene expression by binding to H3K27ac. Previous studies suggest that Brd4 in the NAc is involved in cocaine addiction (Guo et al., 2020), and a recently published paper demonstrates that RVX-208, a domain-selective BET inhibitor, suppresses cocaine-seeking behaviors (Sacko et al., 2026). In addition, Brd4 expression in the prefrontal cortex, hippocampus, and amygdala is accompanied by post-traumatic stress disorder (PTSD)-like behaviors (Wang et al., 2021). Our data imply that *Dkk3* and *Hcrtr2* are candidate targets of Brd4. *Dkk3*, for example, has been reported to be responsive to chronic unpredictable mild stress, and it is associated with depressive and anxiety-like behaviors via the Wnt signaling pathway (X. Chen et al., 2025). It is also reported to be involved in memory maintenance and neuropathic pain (Flores et al., 2024; L. Zhang et al., 2022). Based on these observations, we propose that Brd4 contributes to the downregulation of *Dkk3* and *Hcrtr2* in the NAc after jSI, a hypothesis that remains to be tested by NAc-specific perturbation.

Our analysis also implies that *Dkk3* and *Hcrtr2* are candidate targets of other epigenetic regulators such as Setd1a. Setd1a deficiency causes schizophrenia-like behaviors (Nagahama et al., 2020). Given that human loneliness is associated with the risk of schizophrenia (Liang et al., 2024), and post-weaning SI in mice induces schizophrenia-like behaviors (Pietropaolo et al., 2008), Setd1a may contribute to these behaviors through the regulation of its downstream genes. Besides *Hcrtr2* and *Dkk3*, we found that *Dact1* is another candidate downstream gene that is regulated by Setd1a in jSI mice. *Dact1* encodes a Wnt signaling-associated scaffold protein that regulates dendritic complexity at postsynaptic sites (Arguello et al., 2013; Okerlund et al., 2010), and it is associated with speech and language disorders (Benchek et al., 2021). Based on these observations, we propose that Setd1a activity contributes to the downregulation of these neuronal function-associated genes in the NAc after jSI, a hypothesis that remains to be tested by NAc-specific perturbation.

To summarize, we revealed some changes in gene expression and histone modification levels in the NAc of female mice under jSI stress. Neuronal development-associated and gene expression regulation-associated genes are marked by multiple histone modification alterations near their promoters, and these genes may play an essential role in the response to jSI. Although we did not show the molecular mechanism of histone modification alteration regulating gene expression, we revealed the association between transcriptome and histone modifications in top nodes, hoping to provide directions for future research.

There are also several limitations in this study. First, our DEGs were defined using nominal *p*-values with a small number of biological replicates, and the replicate number also differed between histone modifications; the changes reported here therefore require validation in independent cohorts. Second, we only revealed the association between the transcriptome and multiple histone modifications in NAc neurons. The direct interaction between the regulation of histone modifications and gene expression should be further examined in the future. This could be achieved by manipulating epigenetic factors such as Kdm6b, Brd4, and Setd1a in the NAc or by epigenetic editing via dCas9 system. Third, we inferred candidate epigenetic factors based on previously published public data, selected as the closest available to our system since no dataset in which these factors had been perturbed exists for the NAc. However, different brain regions, cell types, developmental stages, sexes, and experimental conditions between our analyses and public datasets may contribute to the gene expression differences, leading to false negative or false positive results in this study. Fourth, in the NAc, there are different neuronal subtypes such as D1^+^ and D2^+^ neurons, which may have different gene expression and epigenetic regulation patterns. We were unable to separate these subtypes for this current analysis because nuclei were sorted using an anti-NeuN antibody, but future studies are necessary to confirm the roles of these epigenetic regulators in different neuronal subtypes. Finally, this study was performed exclusively in female mice and did not include behavioral testing of the animals used for molecular profiling, so our findings cannot be assumed to apply to males, nor can we relate the molecular changes to behavioral phenotypes in the same individuals.

## Materials and Methods

### Ethics statement

All animals were housed and studied in compliance with protocols approved by the Animal Care and Use Committee of The University of Tokyo. The approval numbers are A2022IQB001 from the Institute for Quantitative Biosciences and 31-7 from the Graduate School of Science. All procedures were followed in accordance with the University of Tokyo guidelines for the care and use of laboratory animals and ARRIVE guidelines.

### Animals and sample collection

Female C57BL/6J mice (CLEA Japan, Inc., Japan) were shipped from the breeder and arrived at the animal facility on postnatal day (PD) 21, at which point isolation was started. Only female mice were used, based on our previous finding that this isolation paradigm produces a behavioral phenotype in females but not males (Sazhina et al., 2025). The juvenile social isolation procedure was performed as described previously (Sazhina et al., 2025). Briefly, mice were randomly assigned to either juvenile social isolation (jSI) or group house (GH) conditions. jSI mice were individually housed in small cages from PD 21 to PD 35. After 2 weeks of jSI, isolated mouse were regrouped with other previously isolated mice and housed in groups of 3-4 per cage. This design allowed us to examine the lasting effects of juvenile isolation rather than the acute effects of ongoing isolation. GH mice were housed in groups of 3–4 per cage throughout the experiment. The animals were housed in a temperature-controlled environment (23 ± 2 °C) with regulated humidity (50 ± 10 %) and were provided *ad libitum* access to food and water. The animals were kept on a 12 h light/dark cycle (lights on at 8 am, off at 8 pm).

In total, 10 control and 9 jSI animals were analyzed. No behavioral testing was performed on these animals, since exposure to behavioral procedures induces transcriptional and epigenetic changes in the brain that would confound those attributable to isolation. Nuclei isolated from NAc of each animal were divided and used for RNA-seq and for all four CUT&Tag experiments, so that all datasets were derived from the same set of animals, and the same number of nuclei was used for every sample in each analysis. No samples were pooled, and each biological replicate corresponds to a single animal. Because a subset of libraries did not pass our quality criteria after sequencing and was excluded, the number of replicates for some datasets is smaller than the number of animals; the replicate number for each dataset is given in the corresponding figure legend.

### Brain tissue collection

Mice at 7 weeks old were deeply anesthetized with isoflurane and euthanized by cervical dislocation. Brains were rapidly removed under ice-cold conditions and sectioned with a brain slicer. The coronal slice containing the NAc was placed on an ice-cold mat (World Precision Instruments), and the NAc was collected using a stainless steel biopsy puncher. Collected tissues were immediately frozen and stored at −80 °C until further use.

### Nuclei isolation

Frozen samples were thawed and kept on ice. Each sample was mixed with 0.5 ml of 54% Percoll buffer (54% Percoll^®^ (Sigma-Aldrich, #P1644), 50 mM Tris-HCl (pH 7.4), 25 mM KCl, 5 mM MgCl_2_, and 250 mM sucrose). Homogenization was performed by pipetting up and down 10 times through a 1 ml syringe fitted with a 23G needle, followed by another 10 repetitions using a 27G needle. Subsequently, 5 μl of 10% NP-40 was added to each sample (final 0.1%), and the tubes were inverted and then incubated on ice for 15 minutes. After incubation, 0.5 ml of 1 × FANS buffer (50 mM Tris-HCl (pH 7.4), 25 mM KCl, 5 mM MgCl_2_, and 250 mM sucrose) was added to each sample using pipette tips pre-coated with 0.5% BSA in PBS. A Percoll density gradient was prepared by sequentially adding 200 μl of 31% Percoll buffer and 100 μl of 35% Percoll buffer at the bottom of the tubes. Samples were centrifuged at 20,000 × g for 10 minutes at 4°C. Debris on the top layer was removed, and 200 μl was carefully collected from the bottom of the tube using BSA-treated tips and transferred to 1.5 ml tubes pre-coated with 0.5% BSA/PBS. For blocking, 100 μl of 7.5% BSA/PBS was added to each sample (final 2.5%), followed by 0.75 μl of anti-NeuN-488 conjugated antibody (Millipore, #MAB377X, 1:400 dilution). After a 10-minute incubation, 50 μl of 31% Percoll buffer was added to the bottom of each tube using 0.5% BSA/PBS pre-coated tips. Samples were centrifuged at 1,000 × g for 10 minutes at 4°C, and the supernatant was discarded. The nuclei pellets were resuspended in 200 μl of 0.2% BSA/PBS and transferred to FACS sorting tubes. Nuclei were sorted using a FACS Melody (Becton Dickinson) with a 488 nm laser and collected into 500 μl of STEM CELLBANKER (Takara Bio). Sorted nuclei were stored at −80°C until further use.

### RNA-seq

Frozen nuclei were thawed at room temperature and centrifuged at 1,000 × g for 5 minutes at 4°C. The supernatant was discarded, and the nuclei were centrifuged again under the same condition. After removing the supernatant, nuclei were resuspended in 3.5 μl of water and transferred to a new tube for cDNA synthesis and library preparation using the SMART-seq Stranded Kit (Takara, #634447) according to the manufacturer’s protocol. Library concentrations were assessed using the TapeStation system (Agilent), and sequencing was performed on the DNB-seq platform (MGI) to obtain 50-base single-end reads.

### CUT&Tag

Frozen nuclei were thawed and bound to concanavalin A (ConA)-coated magnetic beads (BioMag®Plus Concanavalin A, 10 µL per reaction) for 10–20 min on a rotator at room temperature. After fixation with 0.1% formaldehyde in PBS and washing with Wash buffer (20 mM HEPES (pH 7.5) (Sigma Aldrich), 150mM NaCl, 0.5 mM Spermidine, and 1 × Protease Inhibitor (cOmplete)), primary antibodies (anti-H3K4me1 mouse monoclonal antibody: Thermo Fisher, #MABI0302, 1:100 dilution; anti-H3K4me3 rabbit polyclonal antibody: abcam, #ab8580, 1:100 dilution; anti-H3K27ac rabbit monoclonal antibody, Cell Signaling, #8173S, 1:100 dilution; anti-H3K27me3 rabbit monoclonal antibody, Cell Signaling, #9733S, 1:100 dilution) in Primary antibody buffer (0.05% Digitonin (Wako), 2 mM EDTA (pH 8.0), and 0.1% BSA in Wash buffer) were added, and samples were incubated for around 16 h at 4°C with gentle rotation. On the following day, samples were washed and incubated with 100 µL of Dig-wash buffer (0.05% Digitonin in Wash buffer) supplemented with 1 µL of secondary antibody (Rabbit Anti-Mouse IgG, abcam, ab46540, 1:100; Pig Anti-Rabbit IgG, Rockland Immunochemicals, Inc., 611-201-122, 1:100) for 30 min at room temperature. After washing with Dig-wash buffer, pA–Tn5 transposase (Diagenode, C01070001, 0.4 µL per reaction) was added in 100 µL of Dig-300 buffer (300mM NaCl and 0.01% Digitonin in Wash buffer) and incubated for 1 h at room temperature with rotation. Then tagmentation was initiated by adding 100 µL of Dig-300 buffer with 1 mM MgCl₂. Samples were incubated at 37 °C for 1 h. The reaction was stopped by adding 1 µL of 10% SDS and 3.3 µL of 0.5 M EDTA per sample, followed by incubation at 60–65 °C for de-crosslinking for around 16 h. Samples were treated with 1.7 µL of 10 mg/mL Proteinase K (Nakalai) for 30 min at 50 °C. DNA was purified using phenol–chloroform extraction and ethanol precipitation. DNA pellets were washed with 70% ethanol, and the libraries were generated using Q5 Hot Start High-Fidelity 2× Master Mix (New England Biolabs) with the PCR reaction as follows: 5 min at 72 °C and 30 s at 98 °C, followed by 12 cycles of 10 s at 98 °C and 20 s at 63 °C, with a final extension at 72 °C for 1 min. The PCR products were purified with SPRI beads. The purified libraries were quantified using the TapeStation system (Agilent), and sequencing was performed on the DNB-seq platform (MGI) to obtain 100-base paired-end reads.

### Data analysis

#### RNA-seq data analysis

Fastq files were checked to make sure the item “Per base sequence quality” was more than 30 across all positions in read (bp) by FastQC software (http://www.bioinformatics.babraham.ac.uk/projects/fastqc). Trimming was performed by FASTX-Toolkit (http://hannonlab.cshl.edu/fastx_toolkit) and Cutadapt (Martin, 2011) (https://cutadapt.readthedocs.io/en/stable) for files with failed “Adaptor Content” in FastQC Report, and trimmed sequences were no longer than 50 bp. Sequencing reads obtained by HISAT2 were mapped to the reference mouse genome (mm10). Blacklisted regions were removed according to the Encyclopedia of DNA Elements project (Amemiya et al., 2019; Dunham et al., 2012). Counting reads were generated by featureCounts (Liao et al., 2014). edgeR was used to determine the differentially counted reads (Y. Chen et al., 2025). *p*-value < 0.05 and Fold Change > 1.2 were used as the threshold for identifying DEGs. GO analysis was performed by DAVID Bioinformatics (Huang et al., 2007) (https://davidbioinformatics.nih.gov/tools.jsp). Transcription Factor analysis was performed by ChIP-Atlas: Enrichment Analysis (Zou et al., 2024) (https://chip-atlas.org/enrichment_analysis), and “Threshold for significance” and “Distance range from TSS” were set to default. PPI analysis was performed on STRING (Szklarczyk et al., 2023) (https://string-db.org), and the settings were set to default.

#### CUT&Tag data analysis

Quality control, trimming, blacklist removal, and GO analysis were performed as described for RNA-seq. After quality check and trimming, sequencing reads obtained by Bowtie2 (Langmead & Salzberg, 2012) were mapped to the reference mouse genome (mm10), and blacklisted regions were removed. BAMscale was used for counting reads with 5 kbp bin (Pongor et al., 2020), and edgeR was used to determine the differentially counted reads (Y. Chen et al., 2025). *p*-value < 0.05 and Fold Change > 2 were selected as the standard for identifying differentially deposited regions (DDRs), and these DDRs were annotated by the closest gene loci via bedtools with the function of “closest” (https://bedtools.readthedocs.io/en/latest). GO analysis was then performed.

#### Public data analysis

Fastq files were accessed by SRA Toolkit (https://hpc.nih.gov/apps/sratoolkit.html) through the DDBJ database (PRJDB10118 for Setd1a^+/-^) and GEO database (GSE212440 for Kdm6b cKO; GSE63809 for Brd4; GSE123652 for Setd1a^+/-^; GSE181024 for Foxp2^+^ Setd1a^+/-^). Other processes were the same as in “RNA-seq data analysis”.

#### Statistical analysis

The *p*-values of RNA-seq and CUT&Tag were determined from edgeR, and those of GO terms were from DAVID. Fisher’s exact test was used to analyze the association between jSI-induced DEGs and histone modification alterations, as well as epigenetic factors-induced DEGs.

## Supporting information

Revision plan

## Use of AI Tools for Language Editing

GPT-5.4 (OpenAI), Claude Sonnet 4.6, and Opus 5 (Anthropic) were used solely to assist with language translation and editing during the preparation of the manuscript. The following prompts were used:

– “Please translate the following Japanese text into English suitable for an academic paper.”
– “Please proofread the following text to conform to academic writing conventions.”

## Acknowledgments

We thank Etsuko Ogawara, Izumi Yamaguchi, and Tatiana Sazhina (The University of Tokyo) for technical assistance; and members of Kishi laboratories for helpful discussion.

## Author Contributions

**Junpeng You**: Conceptualization, Data Curation, Formal Analysis, Investigation, Methodology, Visualization, Writing – Original Draft Preparation, and Writing – Review & Editing. **Akira Uematsu**: Conceptualization, Investigation, and Resources. **Asuka Jouji-Nishino**: Resources. **Mai Saeki**: Data Curation and Formal Analysis. **Yusuke Kishi**: Conceptualization, Data Curation, Funding Acquisition, Investigation, Methodology, Project Administration, Supervision, Visualization, Writing – Original Draft Preparation, and Writing – Review & Editing.

## Data Availability

The RNA-seq and CUT&Tag dataset was generated from raw sequencing data, and raw datasets (fastq files) or processed datasets have been deposited in the DNA Data Bank of Japan (DDBJ) Sequence Read Archive under the accession code PRJDB42150. The external datasets related to Brd4, Setd1a, and Kdm6b are available in the DDBJ (PRJDB10118 for Setd1a^+/-^) (Nagahama et al., 2020) and GEO database (GSE63809 for Brd4; GSE123652 for Setd1a^+/-^; GSE181024 for Foxp2^+^ Setd1a^+/-^; GSE212440 for Kdm6b cKO) (Chen et al., 2022; Korb et al., 2015; Mukai et al., 2019; Ramesh et al., 2023). Additional data, methods, and codes used in this study are available from the corresponding author upon reasonable request.

---

**Figure S1.**
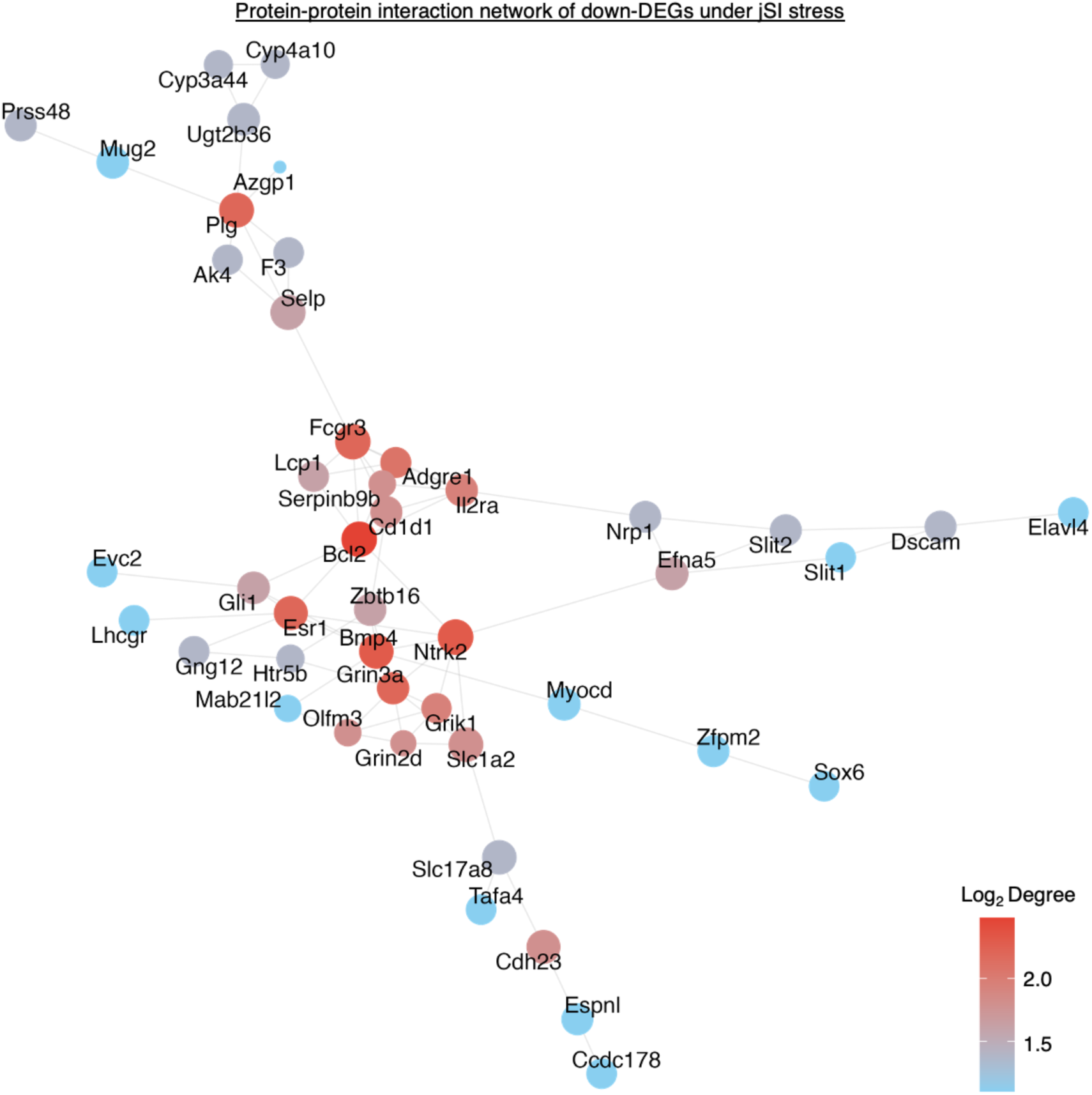
Protein-protein interaction network of down-DEGs under jSI stress.

**Figure S2.**
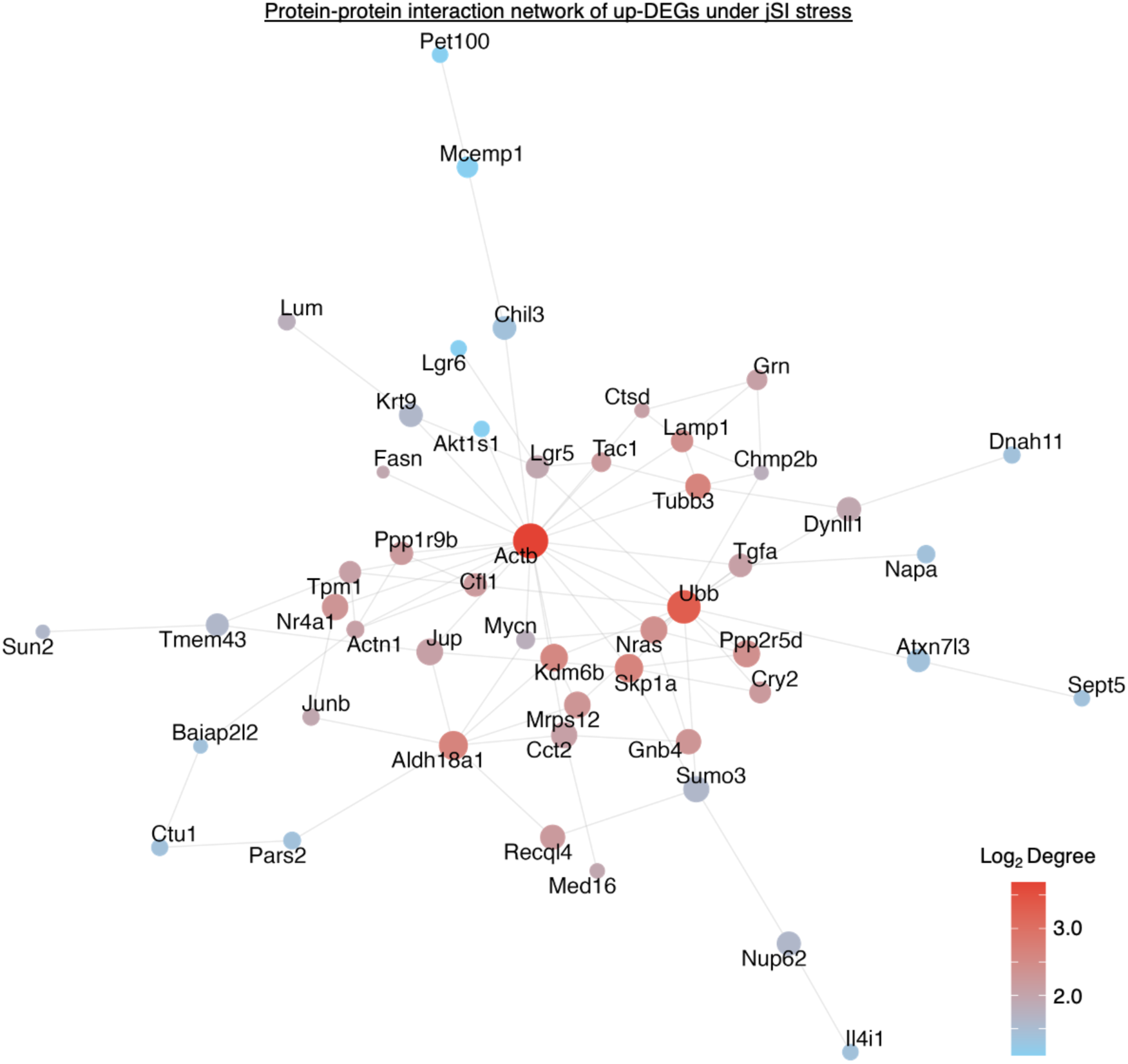
Protein-protein interaction network of up-DEGs under jSI stress.

