## Supplementary material for "Juvenile social isolation induces gene expression alterations and histone modification dysregulations in nucleus accumbens (NAc) neurons of female mice": Revision plan

Review COMMONS

**Manuscript number:** RC-2026-03654
**Corresponding author(s):** Yusuke, Kishi

#### 1. General Statements [optional]

Reviewer #1 (Evidence, reproducibility and clarity (Required))
The manuscript by You et al. investigates changes in gene expression and histone modifications after juvenile social isolation (jSI) in the nucleus accumbens (NAc). They find many differentially expressed genes and find overlap with activating or repressive histone marks. They then go on to compare their data to other published datasets to support their findings. This is an interesting study, and the authors use creative approaches using unique and published data to identify epigenetic mechanisms underlying the changes in gene expression within the NAc. However, there are many points of clarification that are needed to fully evaluate the manuscript and several experimental details.

Reviewer #1 (Significance (Required))
This is an interesting study and the authors use creative approaches using unique and published data to identify epigenetic mechanisms underlying the changes in gene expression within the NAc. However there are many points of clarification that are needed to fully evaluate the manuscript and several experimental details are missing or unclear.

We thank the reviewer for the positive assessment of our study and for the constructive comments, which have helped us to improve the manuscript considerably. The reviewer's points centered in particular on the framing of the Introduction, especially the conflation of adult and adolescent social isolation, the use of nominal rather than FDR-corrected p-values to define DEGs, and the lack of clarity in several parts of the Methods.

We have addressed each of these points in our responses below, in part through revisions already made to the manuscript. Briefly, the Introduction and the Methods have been substantially revised, and we will carry out additional threshold-free analyses to support our conclusions. We are grateful to the reviewer for identifying the points that most needed clarification.

Reviewer #2 (Evidence, reproducibility and clarity (Required))
This paper profiles NeuN positive NAc nuclei after juvenile social isolation. The authors report RNA seq changes and CUT and Tag for H3K4me1, H3K4me3, H3K27ac, and H3K27me3. They then link DEGs to public datasets for Kdm6b, Brd4, and Setd1a. The neuron enriched design is useful. The main claim remains correlative. Causal support is thin.

Reviewer #2 (Significance (Required))

### Revision Plan

Review COMMONS

This study provides a useful neuron-enriched transcriptomic and histone modification resource from the nucleus accumbens following juvenile social isolation. The integration of RNA-seq and CUT&Tag data adds value for researchers studying epigenetic regulation and stress-related neurobiology. However, the advance is primarily descriptive rather than mechanistic, as the conclusions rely largely on correlative analyses without functional validation. The manuscript will be of interest to the neuroepigenetics and psychiatric neuroscience communities, but the conceptual advance is incremental, and the mechanistic claims should be moderated.

We thank the reviewer for recognizing the value of our neuron-enriched dataset and for the constructive comments, which have helped us to improve the manuscript considerably. The reviewer's points centered in particular on the correlative nature of our findings and the need to moderate causal language, the permissive thresholds used to define DEGs and the absence of correction for multiple testing, and the interpretation of the comparisons with published datasets.

We address each of these points in detail in our responses below, and have already implemented a substantial part of the revisions. Briefly, we have removed wording implying that the identified epigenetic factors mediate jSI-induced transcriptional changes and now describe these relationships as associations that remain to be tested, and the comparisons with published datasets are explicitly framed as exploratory. We will also carry out threshold-free analyses and apply correction for multiple testing. We are grateful to the reviewer for these suggestions, which we believe have made the manuscript more accurate about what our data can and cannot support.

Reviewer #3 (Evidence, reproducibility and clarity (Required))

Summary:

In this study, the authors were investigating the effect of juvenile social isolation (jSI) on the nucleus accumbens (NAc) transcriptome in female mice. They used P21 wild type (C57BL6) female, isolated at P21 or group housed, and reunited at P35 to generate their samples for RNAseq on NAc punches. They also performed FACS sorting on the nuclei from NAc lysates to select only neuronal nuclei (NeuN staining). They studied the differentially expressed genes (DEGs) between group housed and jSI by RNAseq. Then, they studied the protein/protein interaction via stringDB on the DEGs identified (up or down) and perform GO analysis on them. They identified Ntrk2, Grin3a, Grik1 and Bcl2; associated with the neuronal function or transcription regulation terms. They also studied the histones modifications (H3K4me1, H3K4me3, H3K27ac, and H3K27me3) after jSI and identified neuronal function and transcription regulation terms again on the DDR (Cut and Tag method). They found that these histones modifications could play a role in jSI-induced adaptations and neuronal function. Finally, they reanalyzed public datasets of RNAseq data to identify histone modifications associated with their DEGs of interest, and compare their DEGs to differences between genotypes in the public datasets (in conditional KO models of some genes of interest, as Kdm6b cKO, BET inhibitor, or Setd1a +/- mice with 3 different mutations. They conclude that histone modification could be involved in jSI-induced gene expression alteration.

### Revision Plan

Review COMMONS

Reviewer #3 (Significance (Required))
Globally, this study is showing transcriptomic and histone modifications occurring after juvenile social isolation in female mice. The authors identified differentially expressed genes linked to neuronal development, regulation of transcription and chromatin remodeling. The reanalyzed public datasets to identify histone modification on the DEG identified and observed the impact of some already published mutations on the gene expression to compare it to their data. The limits of this issue are caused by the reanalysis parts, since the datasets used are cortical and cerebellum samples, in diverse development stage (embryonic, early juvenile, adults - both sexes) that is quite different compared to their paradigm (juvenile females). They also never display the name of the principal DEGs identified (text or plots) which leads to difficult understanding of the findings of this paper. A more focused analysis on NAc or striatal, female only, juvenile stage datasets would be more helpful in this situation. The text should be more precise sometimes and a specific explanation on the exclusion of male mice should be introduce early in the methods.

This study finds its place in the current research on the role of NAc function/dysfunction in behavioral abnormalities induced by social isolation and try to understand the mechanisms behind the abnormal behaviors induced by separation. The audience could be composed of researchers from several domains where social isolation is the cause or the consequence of pathological behaviors, including studies on loss, depression, ASD, Alzheimer, Schizophrenia, etc). The context is quite broad. The results from this paper could help find new molecular targets to alleviate the effects of social isolation and perhaps ameliorate the behavior for mouse models of several diseases or later in patients. A better understanding of the effects of social isolation in female is interesting, but being able to compare both sexes would be even better: identifying sex-differences and common defect is of great interest nowadays in several domains.

The present reviewer has expertise in behavior in mice (both male and female) from juvenile to adult stages, has studied neuronal circuits including prefrontal cortex and striatum (mainly NAc) in behavioral abnormalities in mice in a model of ASD and more recently in an addiction model. The reviewer is interested particularly in sex-differences in neuronal circuits defects and behavior expression in diseases. Finally, the reviewer has recently focused on spatial transcriptomic approaches in addiction models.

We thank the reviewer for the detailed and constructive assessment of our study, and for the many specific suggestions, which have helped us to improve the manuscript considerably. The reviewer's points centered in particular on the discrepancy between the published datasets we reanalyzed and our own experimental paradigm, the absence of gene names in the text and figures, which made our findings difficult to follow, and the need for greater precision in the text, including an explicit explanation of why only female mice were used.

We address each of these points in detail in our responses below, and have already implemented a substantial part of the revisions. Briefly, the rationale for using female mice is now stated in both the Introduction and the Methods, the text has been revised throughout for precision, and the comparisons with published datasets are explicitly framed as exploratory. We will also reanalyze the striatal dataset from Chen et al. (***Sci. Adv.***, 2020), which is considerably closer to our samples than the prefrontal cortex datasets, and we will label the relevant genes in the text and figures. We are grateful to the reviewer for the care taken in reading the manuscript.

#### 2. Description of the planned revisions

#1-2

The authors cite several studies from the Nestler lab on early life stress that link early life stress to histone modifications but failed to cite the manuscripts that investigated the transcriptional changes in response to jSI. These studies also highlight sex differences in jSI. This is important given that this study only uses females. Many of the effects described might not be comparable simply because of the sex of the animals.

We thank the reviewer for this helpful comment. We agree that studies investigating transcriptional changes in response to jSI, including those reporting sex differences, should be cited, and we will add these references in the revised Introduction.

The paragraph highlighted by the reviewer (the third paragraph of the Introduction) was intended to summarize the relationship between stress and epigenetic regulation in the NAc, which is why studies from the Nestler laboratory are prominently represented; transcriptional responses to social isolation in the NAc were summarized in the preceding paragraph. As the reviewer notes, however, other important studies have since been reported, and we will incorporate them accordingly.

We also thank the reviewer for raising the issue of sex differences, which has been highlighted repeatedly by the other reviewers as well and is indeed an important point. We have revised the Introduction to specify the sex of the animals used in the rodent studies, and to make the reported sex differences explicit (lines 75–78).

#1-3

Can the authors clarify if they used an adjusted p-value or nominal p-value. If they are using a nominal p-value the authors should explain their reasoning and provide information regarding if any of the transcripts survived a p-value correction. The addition of threshold-free approaches are more appropriate (GSEA) rather than focusing on transcripts with a nominal p-value. If they are going to present data using a nominal p-value, this should be justified and the cut off should be explained and every interpretation should include a caveat.

We thank the reviewer for this important comment. The reviewer is correct that nominal *p*-values, rather than adjusted *p*-values (FDR), were used in this study, and we agree that this point requires both clarification and revision.

In the revised manuscript, we will take the following steps. First, as the reviewer suggests, we will perform GSEA as a threshold-free approach and replace the current GO analysis of DEGs with the GSEA results. Second, for analyses that necessarily require a defined gene set, namely the PPI analysis, the ChIP-Atlas analysis, and the overlap analyses shown as Venn diagrams, we will continue to use gene sets defined by nominal *p*-values, but we will state explicitly in the text that these analyses are exploratory and hypothesis-generating rather than confirmatory.

Regarding our original choice of threshold, we note that the number of biological replicates in this study (n = 4–6) is small relative to the number of genes tested, and that transcriptional responses to stress in the nervous system are typically of small effect size with substantial inter-individual variability. Under these conditions, FDR correction is highly conservative, and previous studies in this field have reported findings based on nominal *p*-values (e.g. Torres-Berrío et al., ***Nat. Neurosci.***, 2024). Since the aim of the present study was to generate hypotheses rather than to establish definitive causal factors, we adopted a permissive threshold. We will make this rationale explicit in the revised Methods and Results.

Finally, we will add a statement to the Limitations section noting that the findings reported here require validation in future work. We hope that these revisions adequately address the reviewer's concern.

#1-4

It is unclear how the authors confirmed that input RNA or neurons were similar across samples for the library prep. This is especially important given the top genes that are differentially expression. The finding that beta actin (Actb) is up in jSI vs GH animals. The results could be due to differences in input rather than actual differences in expression.

We thank the reviewer for raising this important point. We understand the concern to be that the observed change in *Actb*, a gene commonly regarded as a housekeeping gene, may reflect differences in input material between samples rather than a genuine difference in expression.

We would like to clarify that the same number of sorted nuclei was used for every sample, and we have stated this explicitly in the Methods section of the revised manuscript. Consistent with this, library quality assessed by fluorometry (Qubit) and capillary electrophoresis (TapeStation) was comparable across all samples, and the FACS profiles showed the similar pattern in every case; we will present these data in the point-by-point response accompanying the revised manuscript. In addition, we will report the TMM normalization factors calculated in edgeR, together with PCA and/or MDS analyses, to confirm that no sample deviated substantially from the others and that normalization was applied appropriately.

Regarding housekeeping gene expression, *Actb* is known to undergo activity-dependent changes in expression in neurons and to contribute to synaptic function and plasticity. We consider this an interesting observation in its own right, particularly as another reviewer noted that the “actin reorganization” GO term among our differentially expressed genes warrants further description. In the revised manuscript, we will cite the relevant literature and discuss this point. We will also confirm that the expression of other housekeeping genes was largely unchanged, supporting the consistency of RNA-seq quality across samples.

#1-5

There are aspects of the methods are difficult to understand. For example, under RNA-seq, the authors mention "frozen nuclei were thawed and centrifuged.....the supernatant was discarded and nuclei were centrifuged again under the same conditions" Can the authors clarify what was done here? Were the nuclei resuspended in STEM CellBANKER or something else? While this is a concrete example, there are many other places in the methods the are like this, meaning that steps seem to be skipped and it then becomes difficult to assess the approach. It is recommended that the authors work to clarify the methods. Another example, how the DNA was treated in the CUT&TAG and how much DNA was added to the library prep.

We apologize for the lack of clarity in the Methods section. We will review and revise the entire Methods section to ensure that each step is described unambiguously.

Regarding the specific example raised by the reviewer, sorted nuclei were resuspended in STEM CELLBANKER and frozen in this solution; after thawing, they were centrifuged directly without resuspension in any other buffer. We will make this explicit in the revised text.

For the CUT&Tag experiments, the same number of nuclei was used for every sample, and we have stated this explicitly in the revised Methods. Regarding the amount of DNA used for library preparation, we did not quantify the DNA prior to PCR amplification, as the yield at this step is below the range of reliable quantification and measurement is not included in the original CUT&Tag protocol. Instead, all libraries were amplified with the same number of PCR cycles (12 cycles, as stated in the Methods), ensuring that they were prepared under identical conditions throughout.

#2-major-2

DEG thresholds are loose. DEGs are defined as "p-value < 0.05 and Fold Change (FC) > 1.2." There is no clear FDR cutoff. With ~1250 DEGs from n = 5 to 6, many hits may be noise. Please report FDR filtered lists or justify the uncorrected p value choice. Re run key GO and overlap tests on a stricter set.

We thank the reviewer for this important comment, and we agree that the thresholds used to define DEGs require clarification and revision.

In the revised manuscript, we will take the following steps. First, we will perform GSEA as a threshold-free approach and replace the current GO analysis of DEGs with the GSEA results, so that our functional conclusions do not depend on an arbitrary cutoff. Second, for analyses that necessarily require a defined gene set, namely the PPI analysis, the ChIP-Atlas analysis, and the overlap analyses shown as Venn diagrams, we will continue to use gene sets defined by nominal *p*-values, but we will state explicitly in the text that these analyses are exploratory and hypothesis-generating rather than confirmatory.

Regarding our original choice of threshold, the number of biological replicates in this study (n = 4–6) is small relative to the number of genes tested, and transcriptional responses to stress in the nervous system are typically of small effect size with substantial inter-individual variability. Under these conditions, FDR correction is highly conservative, and previous studies in this field have reported findings based on nominal p-values (e.g. Torres-Berrío et al., ***Nat. Neurosci.***, 2024). Since the aim of the present study was to generate hypotheses rather than to establish definitive causal factors, we adopted a permissive threshold. We will make this rationale explicit in the revised Methods and Results, and we will add a statement to the Limitations section noting that the findings reported here require validation in future work.

#2-major-3
Public data overlaps are hard to interpret. Kdm6b data are from cerebellum. Brd4 data are from cultured cortical neurons treated with JQ1. Setd1a data are mostly PFC. The authors note that "different brain regions, cell types, and experimental conditions... may contribute to... false negative or false positive results." That caveat is important. Overlaps should be framed as hypothesis generating only. Do not treat them as evidence that these enzymes act in NAc under jSI.

We fully agree with the reviewer on this point, and we have revised the manuscript accordingly.

First, we have explicitly framed all comparisons with published datasets as exploratory and hypothesis-generating, and we have removed any wording implying that these enzymes act as mediators of jSI-induced transcriptional changes in the NAc.

Second, as pointed out by Reviewer #3, the study by Chen et al. (***Sci. Adv.,*** 2022) also performed scRNA-seq using the striatum of Setd1a heterozygous mice. Since striatal tissue is far closer to our NAc samples than the prefrontal cortex, we will reanalyze the striatal dataset and compare it with our jSI DEGs. Depending on the outcome of this analysis, we will reorganize the figures so that the most relevant comparison is presented in the main text and the remaining reanalyses are moved to the supplementary material.

Third, we have explained our rationale for dataset selection in the revised text. The three factors examined here were nominated by our own data: Brd4 and Setd1a emerged from the ChIP-Atlas promoter analysis, and Kdm6b was itself upregulated in our RNA-seq data. We then searched for publicly available RNA-seq datasets in which these factors had been perturbed in the nervous system. No such dataset exists for the NAc or striatum for Kdm6b or Brd4, and we therefore selected the datasets that were closest to our system among those available. We have stated this limitation in the Limitation section, together with the differences in brain region, cell type, developmental stage, and sex between these datasets and our own.

#2-major-5
CUT and Tag analysis is coarse for promoter claims. Signals are quantified in "all 5 kbp bins." That bin size can blur promoters, enhancers, and neighboring genes. Please add peak calling or TSS centered analyses for key loci such as Grik1, Bcl2, and Dkk3. Also show more browser tracks beyond one example.

We thank the reviewer for this comment, and we agree that quantification in 5 kbp bins is too coarse to support claims about promoter-level regulation.

In the revised manuscript, we will perform higher-resolution analyses centered on transcription start sites for the key loci highlighted by the reviewer, including *Grik1*, *Bcl2*, and

*Dkk3*, so that promoter signals can be evaluated separately from those of surrounding regions. We will also try to perform peak calling to define regions of enrichment more precisely.

We will also increase the number of browser tracks shown, so that examples are provided for each histone modification rather than for a single locus.

#2-major-6

Multiple testing for overlaps needs attention. Many Fisher tests compare DEGs with DDRs and with several public DEG lists. Report whether p values were corrected across tests. Some reported overlaps are small in absolute numbers even when p values look significant.

We thank the reviewer for this comment. The reviewer is correct that we did not correct for multiple testing across the repeated Fisher's exact tests.

In the revised manuscript, we will apply the Benjamini-Hochberg procedure and report adjusted *p*-values in the figures. Correction will be performed within each analysis category rather than across all tests, namely the overlaps between DEGs and DDRs, and the overlaps between our jSI DEGs and each of the published datasets for Kdm6b, Brd4, and Setd1a. We note that these tests are not fully independent, since they share the jSI DEG list and involve mutually exclusive up- and down-regulated gene sets; the Benjamini-Hochberg procedure remains valid under such positive dependence.

We also agree that *p*-values alone can be misleading when the absolute number of overlapping genes is small. For every comparison, we will additionally report the observed and expected numbers of overlapping genes together with the odds ratio or fold enrichment, so that the magnitude of each overlap can be assessed independently of its *p*-value.

#2-minor-2

Down DEG GO terms include "Chondrocyte differentiation" and "Positive regulation of cartilage development." These look odd for NAc neurons. Check annotation quality and whether these terms survive stricter DEG filters.

We thank the reviewer for this observation. Genes involved in developmental processes are frequently shared across tissues, and GO annotation assigns such pleiotropic genes to multiple terms, which can produce enrichment for categories that appear unrelated to the tissue under study.

In our case, the genes driving the enrichment of "chondrocyte differentiation" and "positive regulation of cartilage development" include *Sox5*, *Bmp4*, and *Nfib*, all of which have established roles in nervous system development. Whether these genes are functionally important in the NAc remains unknown, but their appearance in these terms reflects annotation overlap between chondrocyte and neural developmental programs rather than an implausible result.

We nevertheless acknowledge the concern regarding our DEG thresholds, as discussed in our response to the Reviewer #2's comment (#2-major-2). We will perform GSEA as a threshold-free approach and use it to confirm the functional categories identified by the current GO analysis.

Review COMMONS

#3-major-5（OPTIONAL）

Datasets selection: The public datasets used through this study are far from the original experimental design proposed in this study (cerebellum at P14, cortex at E16.5, nonspecific inhibitor, whole adult PFC = 12-14weeks old). It is important to note that the dataset used, from Chen et al 2022 (whole PFC) also studied the striatum of heterozygous Setd1a mice in the same paper. Why did the author reanalyzed PFC data instead of striatum, which would probably look more like their NAc-restricted samples? Restrict the analysis of the public dataset on NAc data, if possible on females only, and/or try to obtain data from similar experimental design (social isolation, stressed mice). Note here, that the development stage during which the mice have been isolated/regrouped and sample taken will probably of importance. The reanalyzes mix embryonic, early juvenile and adult samples, none of which is consistent nor look like their set up (P31-35).

We thank the reviewer for this thoughtful comment, and we agree that the discrepancy between the published datasets and our own experimental design is a substantial limitation.

Regarding the Chen et al. (***Sci. Adv.***, 2022) dataset, we are grateful to the reviewer for pointing out that the same study also profiled the striatum. Since the NAc is part of the striatum, this dataset is considerably closer to our samples than the prefrontal cortex, and we will reanalyze it and compare it with our jSI DEGs in the revised manuscript.

We will also explain our rationale for dataset selection explicitly. The three factors examined here were nominated by our own data: Brd4 and Setd1a emerged from the ChIP-Atlas promoter analysis, and Kdm6b was itself upregulated in our RNA-seq data. We then searched for publicly available RNA-seq datasets in which these factors had been perturbed in the nervous system. To our knowledge, no dataset combines perturbation of these enzymes with the NAc or striatum (except for the Setd1a striatal dataset noted above), with female animals only, or with a social isolation or stress paradigm. Among the datasets available, we therefore selected those closest to our system, prioritizing perturbation of the factor of interest, since this was the specific question the analysis was designed to address. We will state this constraint clearly in the revised text, together with the differences in brain region, cell type, developmental stage, and sex between these datasets and our own.

Finally, in line with the comments from this reviewer and from Reviewer #2 (#2-major-3), we will frame all of these comparisons as exploratory and hypothesis-generating, and will remove wording implying that these enzymes mediate jSI-induced transcriptional changes.

#3-major-7

Assumptions are made based on 2 sets of reanalyses. These parts should be displayed in the supplementary to help the authors target some genes of interest rather than the principal figures. These analyses didn't seem convincing due to too much shift from the original issue of the paper (which is jSI in the NAc in female mice).

We thank the reviewer for this comment, and we agree that the reanalyses of published datasets are exploratory in nature and are considerably removed from the central question of this study.

As described in our response to the Reviewer #3's comment (#3-major-5), we will reanalyze the striatal dataset from Chen et al. (***Sci. Adv.***, 2022), which is far closer to our NAc samples than the prefrontal cortex datasets used previously. Depending on the outcome of this analysis, we will reorganize the figures so that the most relevant comparison is retained in the main text and the remaining reanalyses are moved to the supplementary material.

Throughout the revised manuscript, we will present these comparisons explicitly as a means of narrowing down candidate genes for future investigation, rather than as evidence that these enzymes act in the NAc under jSI. We have also added a statement at the beginning of this section noting that the datasets were obtained under conditions different from ours, and the specific differences will be described in the Limitations section.

#3-major-8
Volcano plots throughout the study: Should display the genes names (at least top 10 up and top 10 down DEGs) on the graph, otherwise the volcano plots are unreadable.

Thank you for your comment. We will present our top annotated genes in the figure.

#3-minor-intro-10
General comment: Since the paper is focused on female, it could be of interest to state in the introduction if/how sex differences exist in relation to social isolation and human diseases showing isolation as a phenotype.

We thank the reviewer for this suggestion, which we have adopted. We have added a statement to the Introduction describing what is known about sex differences in the effects of jSI in rodents, noting that while some outcomes are shared between sexes, others such as sociability and aggression differ. We will also describe what is known about sex differences in the human conditions in which isolation or loneliness features as a phenotype. We have also specified the sex of the animals used in the rodent studies we cite, as requested by Reviewer #1 (#1-2). We have kept this description focused on social isolation rather than surveying sex differences in psychiatric disease more broadly, so that it remains relevant to the present study.

#3-minor-results-1
Section 1: Only 1 or 2 GO terms (down / up DEGs) are described, but top5 is represented in the figure. Description of the others would be of interest (for example actin reorganization could be particularly interesting). Also, citing some DEGs from the top UP and DOWN, representative of the GO terms, could be of huge interest here.

We thank the reviewer for this comment. We agree that describing the GO terms in more detail would improve the readability of this section. In the revised manuscript, we will describe all of the top-ranked GO terms shown in the figure, rather than only one or two, including terms such as actin reorganization. We will also name representative differentially expressed genes belonging to these terms in the text, so that the reader can appreciate which genes underlie each enrichment without consulting the supplementary tables.

#3-minor-results-2
Figure 1, E/F/G: Some genes in Fig1G are not from top10 nodes in DEGs (Slc17a8, Hcrtr2, Dkk3, Dact1, Nr4a1, Fosl2, Htr5a). They are stated as "potentially important genes" in the legends and are found later in the study as important using other methods than RNAseq. Since Fig1G is displaying RNAseq results, it would be better to stick to the Top10 genes displayed here and put in another figure the other "potentially important genes".

What means "potentially important"? Why these? What are the criterion? Also, the fig1G is unclear visually: separate it in two for top10 down and top10 up, it would be easier to understand and navigate.

Legends: Statistics used for 1G are not stated.

Volcano plot: Top 10 genes UP/down could be displayed on the graph.

We thank the reviewer for these comments.

We agree that including genes in Fig. 1G that were not among the top nodes made the figure difficult to follow. These genes will be moved to the section corresponding to the previous Fig. 5, where they are first identified as candidates, and Fig. 1G will show only the top node genes.

We also agree that the term "potentially important genes" was unclear. Since this panel shows the top nodes identified by the PPI analysis, we have replaced this wording with "top nodes" in the figure legend.

Figure 1G will be divided into separate panels for up-regulated and down-regulated DEGs, as suggested.

We will state the statistical method used in the figure legend, and we will label the top 10 genes on the volcano plot.

#3-minor-results-4
Figure 2: A/B/C: only one mention of the NAc in the data; Fig D: 5/8 NAc datasets. The analysis here seems unbalanced. Why not take into account only NAc datasets? The composition of cortical area or retina is highly different than the NAc (mostly Glutamatergic vs GABAergic populations). If doable, the analysis focused on NAc datasets would be better.

We thank the reviewer for this suggestion, which we agree would improve the specificity of this analysis.

For the histone modification analysis, a sufficient number of NAc-derived datasets is available in ChIP-Atlas to support the analysis on its own, and we will therefore repeat this part using only NAc datasets in the revised manuscript.

For the transcription factor analysis, however, the number of NAc-derived ChIP-seq datasets is too small for a comparable enrichment analysis, and restricting the analysis in this way would leave most candidate factors untested. We will therefore retain the "Neural" category for this part, and we will state explicitly that the underlying datasets derive from a range of neural tissues whose cellular composition differs from that of the NAc, as the reviewer notes. As described in our response to the Reviewer #3's comment (#3-minor-results-3), we will also make clear that this analysis was intended to generate candidates rather than to identify regulatory relationships operating in the NAc.

#3-minor-results-5

Section 3: "First, neuronal development-related genes, such as 'nervous system development', were found in all DDRs of the four histone modifications" line 193-194: sentence is unclear, the author probably meant "term". No gene have been cited here in any histone modification experiment (nor visible in the figure, only dots without names, top 10 up/down could be displayed on the volcano plot). It would be of interest to state at least some of the genes identified here (DDRs, closest loci) and to see if/how many common genes from RNAseq data were found again here. GO terms: again, only one or two examples are described, but figure shows the top5. They all could be at least stated.

The conclusion of the first paragraph states: "These results were consistent with the transcriptome analysis that neuronal function and transcription-related genes were affected, and with the transcription factor analysis that epigenetic regulators were predicted to bind to the promoter regions of these genes." line 197-199: Since the author did not state any genes, we can only believe that the result are consistent based on two vague GO terms "neuronal system development" and "regulation of transcription by RNApol II/chromatin remodeling". If the lector has to read itself every gene table to know which genes are dysregulated in jSI, this study will be really time consuming.

We thank the reviewer for these comments, and we apologize that the description of "nervous system development" was inaccurate. We have rewritten this sentence so that it refers to genes functionally related to this GO term being enriched among the DDRs, rather than to the term itself being found among the DDRs (lines 218–219).

We will also describe all of the top-ranked GO terms shown in the figure, rather than only one or two, and we will name the genes associated with the DDRs both in the text and on the volcano plots. In addition, we will state how many of these genes overlap with the DEGs identified in our RNA-seq analysis, so that the correspondence between the two datasets is apparent without consulting the supplementary tables.

#3-minor-results-6

Figure 3: Volcano could display the top10 names of DDRs.

"The results indicated that down-DEGs were associated with H3K4me1, H3K4me3, and H3K27ac." line 203: In which direction are altered H3K4me1, H3K4me3, and H3K27ac? This is important to know.

"Consistent with their active roles in transcription, downregulation of H3K4me1 and H3K27ac was more relevant to down-DEGs than up-DEGs." line 205: Why? Unclear statement.

"Considering the composite roles of H3K4me3 (an active histone modification) and H3K27me3 (a repressive histone modification), we hypothesize a major role of H3K27me3 in these up-DEGs, and the contribution of H3K4me3 to gene expression alteration by jSI might be small, though we cannot exclude the possibility that it regulates certain genes locally or plays a repressive role." line 208-211: It is very unclear here, why H3K27me3 should play a major role while H3K4me3 alteration "might be small". This has to be further discussed.

"For top 10 nodes among up-DEGs, we didn't find any significant alterations in any of the four histone modifications around their gene loci, except for downregulated H3K4me3 around Aldh18a1, Lamp1, and Gnb4" line 217-219: Formulation is clumsy here, reformulate.

"Some of these genes were marked by multiple altered histone modifications." Line 223: Which ones? Only Bcl2 displayed, but the authors state "some of these genes" right after writing "H3K27ac was found to be downregulated around Grin3a, Grik1, and Adgre1". Are these the other genes showing several histone modifications? It is unclear.

We thank the reviewer for these comments, which have helped us to clarify this section. Regarding the direction of the histone modification changes, we have revised the text to state explicitly in which direction each modification was altered, rather than referring only to an association.

Regarding the roles of H3K4me3 and H3K27me3 in up-DEGs, we agree that our reasoning was not adequately explained. We have rewritten this passage to make the logic explicit: the reduction of the repressive mark H3K27me3 is consistent with the upregulation of these genes, whereas the concurrent reduction of the active mark H3K4me3 is not, which is why we consider H3K27me3 the more likely contributor at these loci (lines 237–241).

We have also rewritten the sentence describing the top 10 nodes among up-DEGs, which was awkwardly constructed, so that it now states positively which genes showed an alteration and in which direction (lines 243–253). Similarly, we have revised the sentence referring to genes marked by multiple altered histone modifications, so that it specifies which gene is being described rather than referring vaguely to "some of these genes".

For the volcano plots, we will label the top-ranked DDRs, as we will also do for the volcano plots elsewhere in the manuscript.

#3-minor-results-7
Figure 4B: only Grik1 as an example. Why only this one and not Bcl2 that moreover show several modifications? Could be helpful to show an example of each modification.

Thank you for your comment. We will present the modification enrichment of specific genes that we mentioned.

#3-minor-results-8
Section 4, Kdm6b: "To examine the possible contribution of Kdm6b to jSI, we re-analyzed the RNA-seq data from Kdm6b-knockout in the published study (Ramesh et al., 2023)." line 240-241: this study is about conditional Kdm6b KO in the cerebellum, on naive P14 male and female mice's neurons in culture. The authors extrapolate the results from a completely different neuronal population/region and sex to justify the potential effect jSI could have on their adolescent female mice. This sentence is misleading for the reader, since the model used (not stressed) and experimental conditions are far from what they are studying. This sentence needs some reformulation to better explain their goal. They show the DEGs (up/down) from reanalyzed data and overlap between these DEGs and the one from Figure1, but the conditions are far from each other here. One could ask what the specificity of their overlap demonstrated here.

"Fosl2 and Nr4a1 are immediate early genes (IEGs) in response to neuronal activation in many brain regions (Dave et al., 2025; Shi et al., 2024), and these two genes have been reported to be involved in memory maintenance (McNulty et al., 2012; Mizuno et al., 2020) and Parkinson's disease (PD) (Fan et al., 2020; Rouillard et al., 2018). In addition, Htr5a, which encodes serotonin receptor 5A, was also upregulated in jSI and downregulated by Kdm6b KO (Fig. 1G, 5G, Table S1). And Htr5a has been reported to be a risk factor of human schizophrenia (Guan et al., 2016)." line 254-260: This part of the result paragraph is about introduction/discussion again. This should be moved appropriately.

We thank the reviewer for these comments.

Regarding the Ramesh et al. (***Elife***, 2023) dataset, we agree that the experimental conditions differ substantially from ours, and that our original wording did not make this clear. We have added a statement at the beginning of this section explaining how the datasets were selected and noting that they were obtained under conditions different from ours, so that the reader understands from the outset that these comparisons were intended to narrow down candidate genes rather than to test whether these enzymes act in the NAc after jSI. As described in our response to the Reviewer #3's comment (#3-major-5), we will also state the specific differences in brain region, cell type, developmental stage, and sex in the Limitations section, and we will reanalyze the striatal dataset from Chen et al. (***Sci. Adv.***, 2022), which is considerably closer to our samples.

Regarding the passage describing *Fosl2*, *Nr4a1*, and *Htr5a*, we agree that the discussion of their roles in memory and disease belongs in the Discussion rather than the Results. We have removed this material from the Results, retaining only the minimal information needed to follow why these genes were of interest, and we will incorporate the remainder into the Discussion. We have applied the same principle to the corresponding passage in the transcription factor section, as described in our response to the Reviewer #3's comment (#3-minor-results-3).

#3-minor-results-10
Section 4, Setd1a: Here, they used 3 separate datasets: whole PFC of Setd1a heterozygous mice (exon 4 LacZ/Neo cassette insertion), whole PFC from loss of function Setd1a heterozygous mice and FoxP2+ nuclei from PFC of Setd1a +/- mice (frameshift in the 15th exon). These datasets are quite different between themselves and compared to NAc samples from jSI mice. The authors stated that the first two datasets had a low DEG overlap with their samples but continued with the third which showed a significant overlap for Hcrtr2, Dkk3 and Dact1.

They finally conclude that: "Taken together, these results suggest that epigenetic factors, such as Kdm6b, Brd4, and Setd1a, may mediate jSI-induced gene expression alterations." Nothing in these datasets is comparable to what they want to prove here, it is a huge stretch to propose these genes as mediators of jSI. Reformulate. These results could be exploited as exploratory, to reduce the number of potential targets, but needs to be investigated on their own.

We thank the reviewer for these comments, with which we largely agree.

Regarding the differences between the three Setd1a datasets and our own samples, we have stated these in the Limitations section (lines 514–519), as described in our response to the Reviewer #3's comment (#3-major-5). We will also reanalyze the striatal dataset from the same study by Chen et al. (***Sci. Adv.***, 2022), which is considerably closer to our NAc samples than the prefrontal cortex datasets, and we will reorganize this section accordingly.

Regarding the difference in overlap between the three datasets, we would note that all three comparisons were performed and reported, and that the low overlap with the Mukai and Nagahama datasets was described in the original manuscript rather than omitted. One possible explanation for this difference is that the Chen dataset was generated from sorted Foxp2-positive nuclei, whereas the other two were derived from whole prefrontal cortex, in which signals from neurons may be diluted by non-neuronal cell types. We have added this to the text as a possible interpretation rather than a demonstrated explanation, and we have stated explicitly that all three datasets were compared in the same way (lines 318–322).

Finally, we agree that proposing these enzymes as mediators of jSI-induced gene expression changes overstates what our data support. We have removed such wording and reframed these results as exploratory analyses that narrow down candidate genes for future investigation (lines 328–332), as described in our response to the Reviewer #2's comment (#2-major-1).

#3-minor-results-11
Figure 5: volcano plots: top10 genes visible could be useful. This section of the results would fit better displayed in the supplementary, since they reanalyzed datasets far from their experimental conditions. These genes of interest should be further investigated in their jSI model.

As described in our response to the Reviewer #3's comment (#3-major-7), we will reorganize this section in light of the reanalysis of the striatal dataset, retaining the most relevant comparison in the main text and moving the remaining reanalyses to the supplementary material. We have also revised the text so that each of these sections concludes by identifying the genes concerned as candidates requiring further investigation in our jSI model, rather than as established targets. We will label the top differentially expressed genes on the volcano plots, as described in our response to the Reviewer #3's comment (#3-minor-results-2).

#3-minor-discussion-7
"Besides these two main shared functions affected by jSI, our results suggest that other biological processes are potentially mediated by one or more histone modifications. For example, some DDRs of H3K27ac and H3K27me3 are functionally enriched around cell adhesion-associated genes, and this is consistent with previous papers suggesting that cell adhesion is affected by isolation (Santiago et al., 2023; Wu et al., 2022)..." line 377-382: This GO term appeared in the figure, but has never been mentioned clearly in the results. The explanation goes on for a full paragraph. It could be better to introduce it before if it is of interest. Also, which cell-adhesion genes have been found in the RNAseq / cut&tag experiments for this family of genes (never stated)?

We thank the reviewer for this comment. We agree that discussing cell adhesion at length in the Discussion is inappropriate when the corresponding GO term was never described in the Results, and that the genes underlying this enrichment were not identified.

We will introduce this GO term in the Results section, where the functional enrichment of the DDRs is described, and we will name the cell adhesion-associated genes identified in our RNA-seq and CUT&Tag analyses both there and in the Discussion. This will be done together with the more comprehensive description of the top-ranked GO terms that we will add in response to the Reviewer #3's comments (#3-minor-results-1 and #3-minor-results-5).

#### 3. Description of the revisions that have already been incorporated in the transferred manuscript

#1-1
The text in the introduction conflates adult and adolescent social isolation which have very different effects on behavior. Additionally, there is evidence that isolation during adolescence can have permanent effects on behavior but the behavioral effects of adult isolation in rodents are transient. It is recommended that the authors restructure the intro to be more specific to describing the adolescent period and why epigenetic mechanisms would be expected to regulate changes induced by jSI.

We thank the reviewer for pointing out that adult and adolescent social isolation are conflated in the Introduction. We agree that the effects of social isolation differ between adulthood and adolescence, and we have revised the Introduction to clearly distinguish between the two, with a specific focus on the adolescent period. Specifically, we now note that isolation during adolescence can produce lasting behavioral alterations, whereas the effects of adult isolation are largely transient, and we have added a statement explaining why the adolescent period may therefore be particularly susceptible to epigenetic reprogramming (lines 69–74). In addition, we now explain why epigenetic regulation is a plausible candidate mechanism for the changes induced by jSI: since the effects of jSI persist long after the isolation period has ended, environmental stress during this window is likely to leave a lasting molecular trace within affected cells, and epigenetic regulation can stably maintain altered transcriptional states (lines 93–96).

#1-6

Can the authors please explain why only females were used for these experiments? In addition, can the authors please comment on potential caveats in the interpretation by only including females in the study?

We thank the reviewer for raising this point, which was also noted by the other reviewers. We apologize that this rationale was not stated explicitly in the original manuscript, and that our previous work was not cited in this context.

Our focus on female mice was not arbitrary but followed from our previous work using the same isolation paradigm as in the present study. In Sazhina et al. (***Neuroimage***, 2025), in which mice were isolated from P21 to P35 and regrouped from P35 to P49, we found that jSI produced a heightened fear response in female but not male mice. Since the NAc has been implicated in scaling fear responses to threat intensity, and since this region undergoes a critical period around P28, we hypothesized that isolation during this window disrupts NAc development in a manner that leads to inappropriate fear responses in adulthood, and that this underlies the female-specific phenotype we had observed. We therefore designed the present study to examine molecular changes in the NAc of female mice. We have stated this rationale explicitly in the Introduction and Methods of the revised manuscript (lines 73–74, 542-543), and added the relevant references.

We also agree that restricting the study to females limits the interpretation of our findings. Because jSI is known to produce sex-dependent effects on both behavior and gene expression, the alterations reported here cannot be assumed to occur in males, and comparisons with published datasets derived from male or mixed-sex animals must be made with this in mind. We have discussed these caveats explicitly in the Limitations section (lines 524–527), noting that a parallel analysis in males would be required to distinguish shared mechanisms of jSI from sex-specific ones.

#1-7

Were females shipped to the facility on P21? It is unclear.

We apologize that this was not clearly described in the original manuscript. Female mice were shipped from the breeder and arrived at our animal facility at P21, at which point isolation was started directly. We have stated this explicitly in the revised Methods section (lines 540–541). Group-housed control animals were shipped and received on the same day and under the same conditions, so that both groups experienced identical transport.

#1-8
Were any animals used for multiple endpoints or was each endpoint a separate cohort? Were any samples pooled?

We apologize for not describing this clearly. Nuclei were isolated from the NAc of a single animal and divided into five aliquots, each of which was used as one sample for RNA-seq or for one of the four CUT&Tag experiments. Each biological replicate therefore corresponds to a single animal, and no samples were pooled; all endpoints were derived from the same set of animals rather than from separate cohorts. Both our RNA-seq and CUT&Tag protocols have been optimized for use with small numbers of nuclei, which allowed all five libraries to be prepared from a single animal. We have stated this explicitly in the revised Methods section (lines 554–563).

#2-major-1
Causality is not shown. The Discussion states: "Although we didn't show the molecular mechanism of histone modification alteration regulating gene expression, we revealed the association between transcriptome and histone modifications." That limit should shape the Abstract and title more clearly. Phrases like "epigenetic alterations may also play a role" are fine. Stronger wording about mediation should be toned down until NAc specific perturbation is done.

We agree with the reviewer that this study is hypothesis-generating and does not demonstrate causality. We were mindful of this in preparing the original manuscript, but we acknowledge that language implying mediation remained in several places. We have gone through the entire manuscript, including the Abstract, and revised the wording so that it accurately reflects the correlative nature of our findings. In particular, we have removed expressions implying that the identified epigenetic factors mediate jSI-induced transcriptional changes, and now describe these relationships as associations that remain to be tested by NAc-specific perturbation (lines 34-35, 40-42, 197-200, 330-332, 433-434, 465-467, 481-483, 493-496).

Regarding the title, we would prefer to retain the current wording. The title states that jSI is accompanied by alterations in gene expression and in histone modifications, and does not assert that the latter mediates the former; we therefore believe it does not overstate our findings. We note that the title has been modified to specify the sex of the animals used, as requested by Reviewer #3.

#2-minor-1

Only female mice were used. State this early and discuss sex limits. Juvenile isolation effects often differ by sex.

We thank the reviewer for this comment, and we agree on both points.

As described in our response to the Reviewer #1's comment (#1-6), our focus on female mice followed from our previous work using the same isolation paradigm, in which jSI produced a heightened fear response in female but not male mice. We have stated this rationale explicitly in the Introduction and at the beginning of the Methods (lines 524-527, 542-543), and we have made clear in the Abstract and the title that this study was performed in female mice (line 42).

We also agree that the sex-specific limitations of our findings require explicit discussion. Since jSI is known to produce sex-dependent effects on both behavior and gene expression, our results cannot be assumed to generalize to males, and comparisons with published datasets derived from male or mixed-sex animals must be interpreted with this in mind. We have addressed these points in the Limitations section of the revised manuscript.

#2-minor-3
Figure 1 lists "II2ra" in the top nodes table. That is likely Il2ra. Please correct.

We thank the reviewer for catching this error. The reviewer is correct that the gene name in the top nodes table should read *Il2ra* rather than "II2ra", and we have corrected this in Figure 1.

#2-minor-4
Sample sizes differ a lot across marks. H3K4me1 and H3K27me3 have n = 4 in jSI. Discuss power and why replicates differ.

We thank the reviewer for this comment. In this study, 10 control and 9 jSI animals were prepared, and all analyses were performed on nuclei derived from each of these animals. However, a subset of CUT&Tag libraries failed to pass our post-sequencing quality criteria and was excluded from the analysis, which resulted in the differing numbers of replicates across histone modifications. We have described this in the revised Methods (line 554), together with the quality criteria used for exclusion, so that the basis for these differences is transparent.

We also agree that the reduced number of replicates for H3K4me1 and H3K27me3 lowers the statistical power for these marks relative to the others, and that the number of differentially distributed regions detected for them may therefore be underestimated. We have stated this explicitly in the Limitations section.

#2-minor-5
The isolation protocol includes regrouping from P35 to P49. Make clear that effects are lasting post isolation effects, not acute isolation effects.

We thank the reviewer for this comment, which correctly identifies our intent. The regrouping period was included precisely because our interest is in the effects of juvenile isolation that persist into adulthood, rather than in the acute consequences of isolation itself. Our previous work using the same protocol demonstrated a lasting fear phenotype in female mice after the regrouping period (Sazhina et al., ***Neuroimage***, 2025), and a central aim of the present study is to ask whether epigenetic regulation contributes to the persistence of such environmentally induced changes.

We have stated this rationale explicitly in the Methods (lines 548–550), and the Introduction now notes that the behavioral effects of jSI persist long after the isolation period has ended (lines 69–72, 93–94).

#2-minor-6
Methods say "GPT-5.4... and Claude Sonnet 4.6... was used." Fix subject verb agreement.

We thank the reviewer for pointing this out. We have corrected the subject-verb agreement in this sentence of the Methods (lines 682–683), which now reads "GPT-5.4 ... and Claude Sonnet 4.6 and Opus 5 ... were used".

#2-minor-7
Data Availability lists "GSE3508789." Confirm this accession. It looks malformed.

We thank the reviewer for catching this. The accession number was indeed malformed; the correct accession is GSE123652, and we have corrected it in the Data Availability section (lines 672, 717).

#2-minor-8
Abstract keywords include "Loneliness." The mouse work is social isolation. Keep that distinction clear, as the Introduction already does.

We thank the reviewer for this comment. We agree that including "loneliness" as a keyword was inappropriate given that this study examines social isolation in mice, and we have removed it from the keyword list (lines 44–45).

#3-major-1
Title: Add the sex: "in female mice" since it is specific.

We agree with the reviewer and have revised the title to specify that this study was performed in female mice (lines 1–3).

#3-major-2
Referencing: Biorender.com has been used to generate some schematics in this study, but it is never acknowledged or referenced.

We thank the reviewer for pointing out this omission. The schematic in Figure 1 was created using BioRender, and we have added the citation to the figure legend in the format specified by BioRender, together with the corresponding publication license (line 737).

#3-major-6
Wording: Formulation throughout the current study is vague, sometimes misleading, with some unclear sentences (see minor comments for the sentences showing problems). This issue needs to be checked again.

Thank you for your comment. The responses could be checked in minor comments part.

#3-minor-intro-1
"These negative effects are further supported by evidence from Covid-19 during the last few years" line 56/57: reformulate.

We thank the reviewer for this comment. We agree that the original sentence was awkwardly phrased, since it referred to evidence "from Covid-19" rather than to the studies conducted during that period, and since "the last few years" was vague. We have rewritten it to state that studies conducted during the COVID-19 pandemic, when social contact was widely restricted, provided further evidence for these negative effects (lines 56–58).

#3-minor-intro-2
"In addition, isolation contributes to severe social issues, such as increased human suicide risk" line 57/58: issues is plural, but only one example is given; moreover, the term "social issue" associated to suicide is poorly-worded.

We thank the reviewer for this comment. We agree that the plural "issues" was not supported by the single example given, and that describing suicide as a "social issue" was poorly worded. We have rewritten the sentence so that social isolation is described as being associated with adverse outcomes, including the risk of suicide (lines 59–60).

#3-minor-intro-3
"In the case of rodents, socially isolated animal models are proposed to be associated with various human diseases" line 59/60: clumsy sentence, the animal models are associated to human disease? This could be reformulated.

We thank the reviewer for this comment. We agree that the original sentence was awkwardly constructed, since it stated that the animal models themselves were associated with human diseases. We have rewritten it so that the socially isolated animals are described as exhibiting a range of behavioral abnormalities, and these abnormalities, rather than the models themselves, are described as modelling aspects of human diseases such as depression and schizophrenia (lines 60–63).

#3-minor-intro-4
"and the effects of juvenile social isolation (jSI) on motor, emotional, learning, and sociability-related behaviors in rodents have been widely reported (Li et al., 2021; Powell & Swerdlow,

2023; Walker et al., 2019), which further provides evidence of the pathogenesis and molecular mechanisms of human mental disorders." line 63-66: Examples of behavioral dysfunctions would be appreciated here.

We thank the reviewer for this suggestion. We have added examples of the behavioral dysfunctions reported after jSI, namely hyperactivity, elevated anxiety, impaired spatial learning, and altered social play (lines 66–68). In the same paragraph we have also added a description of the sex differences reported for these effects, and a reference to our own previous work using the same isolation paradigm, as described in our responses to the Reviewer #1's comments (#1-2 and #1-6).

#3-minor-intro-5
"The nucleus accumbens (NAc) is a critical component of the brain reward circuitry, and dysfunction of the NAc is associated with drug addiction (Zinsmaier et al., 2022), impaired social interaction (Pomrenze et al., 2022; Shan et al., 2022), and abnormal emotion expression (Gebara et al., 2021)" line 67-70: here, the statement reads as dysfunction of the NAc is responsible of abnormal behaviors (addiction, social behavior or emotional expression), but the articles show that NAc is dysregulated in models of these pathologies. Is the dysfunction in the NAc responsible of or a consequence of the pathologies? This could be better formulated.

We thank the reviewer for this comment. We agree that the original wording could be read as asserting that NAc dysfunction causes these conditions, whereas the cited studies show that the NAc is dysregulated in models of them. We have rewritten the sentence so that NAc dysfunction is described as having been reported in animal models of these conditions, without implying a direction of causation (lines 81–85).

#3-minor-intro-6
"An fMRI study showed that activity of the human NAc is associated with the sense of loss (Cooper et al., 2009; O'Connor et al., 2008)." line 70-72: Sentence says one study, but two references are used. The sentence refers to O'Connor only. Cooper is about reward/effort and NAc activity, not grief/loss, reformulate. Also, "activity" is unclear, the authors could be more precise with "hyperactivity of the NAc has been found in people suffering from loss".

We thank the reviewer for pointing out these problems. We have removed the citation to Cooper et al. (2009), which concerns reward and effort rather than grief, so that the sentence now refers only to O'Connor et al. (***Neuroimage***, 2008) (lines 85–86). We have also replaced the vague reference to "activity" with a statement that hyperactivity of the NAc was reported in individuals experiencing loss, as the reviewer suggested.

#3-minor-intro-7
"The NAc from lonely individuals showed key differentially expressed genes (DEGs) that are associated with both neurodegenerative and neuropsychiatric diseases" line 74-75: Unclear,

give examples of the pathologies here to be consistent with the next sentence about female rats (Alzheimer, Parkinson, Huntington).

We thank the reviewer for this suggestion. We have added the specific diseases identified in that study, namely Alzheimer's disease, Parkinson's disease, and major depression disorder, so that this sentence is consistent with the following sentence describing the findings in female rats (lines 88–92).

#3-minor-intro-8
The next paragraph (line 79-95) about DNA methylation and histone modifications is missing a general conclusion: what is interesting or needs to be more studied? Also, H3K9 and H3K79 have been introduced but unused in the paper, while H3K27ac/me3 have not been introduced. What is known about them?

We thank the reviewer for these comments. We agree that this paragraph lacked a conclusion and that the histone modifications discussed did not match those examined in this study.
We have shortened the description of H3K79 methylation and removed the statement concerning H3K9, neither of which is examined here. In their place we have added a description of the four modifications we analyse, namely H3K4me1, H3K4me3, H3K27ac, and H3K27me3, together with what is known about their roles in the NAc (lines 108–118). The paragraph now concludes by noting that little is known about H3K4me3 and H3K27ac in the NAc under stress, and that how any of these modifications are altered after jSI remains unknown, which motivates the present study.

#3-minor-intro-9
"How do epigenetic elements mediate gene expression dysfunction under jSI stress? In this study, we aimed to reveal the alterations in gene expression and histone modifications induced by jSI, and to elucidate their roles in the context of psychiatric disorders promoted by jSI" line 96-99: Statement is too general, it is missing the term "NAc" here.

We thank the reviewer for this comment. We agree that our statement of aims was too general and omitted the brain region under study. We have rewritten both the question that opens this paragraph and the statement of aims so that they specify the NAc, and we have also indicated that the study was performed in female mice (lines 119–122). In addition, we have removed the wording implying that epigenetic elements mediate gene expression dysfunction, in line with the request from Reviewer #2 that causal language be moderated (#2-major-1).

#3-minor-methods-1
Female mice only have been used in this study. It is never explained why so. Knowing that many diseases show sex differences in prevalence or symptom expression, it would have been helpful to include also male mice in the study, to identify common mechanism linked to jSI versus sex-specific alterations.

We thank the reviewer for this comment, and we apologize that our rationale was not stated in the original manuscript. As described in our response to the Reviewer #1's comment (#1-6), our focus on female mice followed from our previous finding that the same isolation paradigm produced a behavioral phenotype in females but not males (Sazhina et al., ***Neuroimage***, 2025). We have added a statement to this effect at the beginning of the Animals and sample collection section (lines 542–543), and we have discussed the resulting limitations in the Limitations section (lines 524–527).

#3-minor-methods-2
Cut & Tag: Dilution of antibodies is not displayed ("1µL") and the reference for antibodies is unclear: "primary antibodies (H3K4me1, MABI, 536 MABI0302; H3K4me3, abcam, ab8580; H3K27me3, CST, 9733S; H3K27ac, CST, 537 8173S; 1 µL per reaction)". This should be adapted to look like the FACS antibody description: "anti-NeuN-488 conjugated antibody (Millipore, #MAB377X, 1:400 dilution)".

We thank the reviewer for pointing this out. We have revised the description of the CUT&Tag antibodies so that it follows the same format as the FACS antibody description, specifying the host species and clonality, the supplier, the catalogue number, and the dilution for each antibody, in place of the volume per reaction given previously (lines 611–615).

#3-minor-methods-3
"Frozen nuclei were thawed and bound to concanavalin A (ConA)-coated magnetic beads (BioMag®Plus Concanavalin A, 10 µL per reaction) for 10-60 min on a rotator at room temperature." Why so much difference in incubation time here?

We thank the reviewer for pointing this out. We have checked our experimental records and confirmed that the incubation was performed for 10–20 min in all experiments reported here, and we have corrected the text accordingly (line 608). The wider range given in the original manuscript reflected our general protocol, in which we have confirmed that binding is satisfactory anywhere between 10 and 60 min, but this was not the range actually used in the present study.

#3-minor-methods-4
Number of animals: While reading the manuscript, it was unclear that 5 different groups of mice have been used. The number of animals should be stated in the methods in the "nucleus extraction" part or "animals" section to facilitate understanding the methods.

We thank the reviewer for this comment. We have renamed this section "Animals and sample collection" and added a paragraph stating the number of animals analysed in each group, that nuclei from each animal were divided between the RNA-seq and the four CUT&Tag experiments, and that no samples were pooled (lines 539, 554–563). We have also explained that the number of replicates is smaller than the number of animals for some datasets because a subset of libraries did not pass our quality criteria, and we note that the replicate number for each dataset is given in the corresponding figure legend.

#3-minor-methods-5
Data analysis: "For RNA-seq data, p-value < 0.05 and Fold Change > 1.2 were used as the threshold for identifying DEGs. For CUT&Tag data, p-value < 0.05 and Fold Change > 2 were selected as the standard for identifying DDRs." Cut&Tag p-value and threshold is written in RNAseq section, move it to its proper part hereafter "CUT&Tag data analysis".

Thank you for your comment. We revised it (lines 664–665).

#3-minor-methods-6
Suggestion DDR: acronym is present in the methods but not explained. It is however explained in the results. This depends on the order in the publication, but if methods appear first, it would be helpful to understand what stands for DDR.

Thank you for your comment. We revised it (line 665).

#3-minor-methods-7
Cut&Tag data analysis: "The procedures of quality check and trimming were the same as RNA-seq data analysis." line 591; "and the removal of blacklisted regions was the same as RNA-seq" line 594; "GO analysis was the same as RNA-seq analysis." line 599: These parts could be ameliorated to avoid repetition. Since the preparation of nuclei and most of the analysis are the same, the methods could be more straightforwardly explained separating common preparation from specific analysis.

Thank you for your comment. We revised it (lines 659–660).

#3-minor-methods-8
Methods explaining how the authors performed the reanalysis of the public datasets is missing.

We thank the reviewer for pointing out this omission. We have added a "Public data analysis" section to the Methods, describing how the raw data were retrieved from the DDBJ and GEO databases, and how they were processed (lines 667–671). Steps shared with our own datasets are indicated as such rather than repeated in full.

#3-minor-methods-9
Animals have been separated at P21 and regrouped at P35: It is not stated if they were regrouped together or with a group of unstressed WT never separated, which could influence their behavior and stress levels.

We thank the reviewer for this comment. We have clarified that the isolated mice were regrouped with other previously isolated animals, rather than with mice that had never been separated (lines 547–550). As described in our response to the Reviewer #2's comment (#2-minor-5), we have also stated why the regrouping period was included.

#3-minor-methods-10
No behavioral test has been performed on these mice. It would have been appreciated to see that 2-weeks social isolation was efficient to generate stress in these animals (anxiety test, sociability at least). And if this protocol has been previously used in their lab, at least to explain briefly what behavior abnormalities the jSI was inducing.

We thank the reviewer for this comment, and we apologize that this information was not included in the original manuscript.

This isolation protocol has been used previously in our laboratory, and the resulting behavioral phenotype was reported in Sazhina et al. (***Neuroimage***, 2025), in which the same paradigm produced a heightened fear response in female mice. We have added this finding to the Introduction (lines 73–74), and we have also cited it in the Methods as the basis for our use of female animals (lines 542–543), so that the behavioral consequences of the paradigm are documented.

As described in our response to the Reviewer #2's comment (#2-major-4), behavioral testing was not performed on the cohorts used for molecular analysis, in order to avoid introducing transcriptional and epigenetic changes unrelated to isolation.

#3-minor-results-3
Section 2: The first paragraph here is about which Transcription Factor (TF) is predicted to participate in their DEG's expression. Half of this paragraph is introduction about the function of several TF. This is not part of results and should be moved appropriately in the introduction or discussion section, or shortened significantly, since it is now longer than the result part. Importantly here, the analysis is done on ChIP Atlas (public datasets).

They state: "Taken together, promoter analysis of DEGs suggests that potential epigenetic mechanisms may act upstream of jSI-induced transcriptional dysregulation in the NAc." line 176, but the database has never been stated to be NAc-only data nor data from jSI animals. If not, this sentence has to be modified. It was unclear globally if this part was based on their work or data mining on a first read, it should be more clearly stated at the beginning that this is exploratory.

We thank the reviewer for these comments.

We agree that the introductory description of the transcription factors was disproportionately long for a Results section. We have shortened it substantially, retaining only the information required to follow why these factors were of interest, namely that Setd1a and Brd4 act through the histone modifications examined in this study (lines 184–190). The remaining background, including the association of these factors with neurological and psychiatric disease, has been moved to the Discussion.

We also agree that the exploratory nature of this analysis was not stated clearly. The ChIP-Atlas database is compiled from published ChIP-seq experiments and does not contain data from the NAc of socially isolated animals. We have stated this at the outset of the section (lines 177–180), so that the reader understands from the beginning that the analysis was intended to generate candidates rather than to identify regulatory relationships operating in our system, and we have revised the concluding sentence so that it no longer implies that these mechanisms were demonstrated in the NAc under jSI (lines 199–200).

#3-minor-results-9
Section 4, Brd4: Here, the authors reanalyzed data from E16.5 cortical neuronal culture treated with or without BET family inhibitor, which they state is not selective of Brd4 (even if it is part of the BET family). The crossover between this embryonic cortical neuronal population treated with nonspecific inhibitor and their model (juvenile Social Isolation, NAc) is a bit of a stretch. What do the overlap in DEGs really mean here?

"We also examined the possible downstream Brd4 target genes within the gene sets of down-DEGs by jSI and down-DEGs by JQ1 treatment, and we identified Hcrtr2 and Dkk3 in these gene sets (Fig. 1G, 5H, Table S1). Hcrtr2 encodes an orexin receptor, and it has been reported to be involved in altered arousal levels through dopamine neurons (Bandarabadi et al., 2024). Dkk3 inhibits Wnt signaling and is reported to be related to anxiety and memory formation (X. Chen et al., 2025; Flores et al., 2024)." line 275-280: What is the conclusion on these results?

We thank the reviewer for these comments.

Regarding the Korb et al. (***Nat. Neurosci.***, 2015) dataset, we have added a statement at the beginning of this section explaining how the datasets were selected and noting that they were obtained under conditions different from ours, so that the reader understands from the outset that these comparisons were intended to narrow down candidate genes rather than to test whether these enzymes act in the NAc after jSI (lines 261–265). As described in our response to the Reviewer #3's comment (#3-major-5), the specific differences are also stated in the Limitations section (lines 514–519).

Regarding the meaning of the overlaps, we agree that our original wording did not convey this clearly, and in particular that opening with "as expected" was misleading given that we also observed a significant overlap in the opposite direction. We have rewritten this passage so that the overlap in the unexpected direction is stated as an independent observation rather than as a qualifying clause, and we now state explicitly that the two gene sets are related but that the direction of change does not correspond in a simple manner (lines 295–301).

Regarding the conclusion of this section, we agree that none was previously given. We have removed the description of the roles of *Hcrtr2* and *Dkk3* in arousal, anxiety, and memory, which belongs in the Discussion, and have instead concluded the section by identifying these two genes as candidates whose expression may be regulated by Brd4 in the context of jSI (lines 304–306). We have applied the same principle to the corresponding passages in the Kdm6b and Setd1a sections, as described in our response to the Reviewer #3's comment (#3-minor-results-8).

#3-minor-discussion-1
" For example, the expression of glutamate receptors is reduced in the NAc, prefrontal cortex, and hippocampus under isolation stress (Hermes et al., 2011; Mao et al., 2022; Sestito et al., 2011)." line 315-317: Which GluR are reduced here? It needs to be more precise for the reader here, and to state if some genes have been found in common between this literature and their DEGs.

We thank the reviewer for this comment. We agree that the original sentence was too vague, and we have revised it to specify which glutamate receptors were reduced in the cited studies (lines 343–351), namely GluA1 and GluA3 in the NAc and caudate putamen under chronic social isolation stress.
We now also state explicitly how these findings relate to our own data. *Gria1* and *Gria3*, encoding GluA1 and GluA3, respectively, were not among our DEGs, but we identified other glutamatergic synapse-associated genes, including *Grin3a* and *Grik1*, and we note that isolation may therefore affect glutamatergic signaling in the NAc through a partly distinct set of genes under our conditions.

#3-minor-discussion-2
"The NAc is a key component of the brain reward circuit, and it is involved in drug addiction and social behavior (Pomrenze et al., 2022; Zinsmaier et al., 2022). NAc neurons receive glutamatergic inputs from the PFC, basolateral amygdala (BLA), hippocampus, and ventral tegmental area (VTA) (Arrondeau et al., 2024; Dieterich et al., 2021; Elam et al., 2025; Le Borgne et al., 2025; Zinsmaier et al., 2022), and neurons in the NAc output the information to the ventral pallidum (VP) (Liu et al., 2022), VTA (Qi et al., 2022), and other areas of the basal ganglia (Lanciego et al., 2012)." line 318-324: These lines are describing the circuitry of the NAc, some of its inputs (no mention of dopamine afferences from the VTA) and outputs. No use of this information is used after, since they conclude the paragraph with: "Thus, deficits in glutamatergic synapses possibly mediate jSI-induced behavioral abnormalities, including impaired social interaction, anxiety, and an increased risk of substance abuse." line 325-326: What is the point of describing the circuit, if it is not interpreted regarding their results? What is their hypothesis on the circuit dysfunction in jSI female mice? They were discussing the DEGs from RNAseq result before this paragraph. What is the link/hypothesis between their DEGs and the glutamatergic circuits of the NAc? Is the NAc directly responsible of jSI-induced behavioral abnormalities for them or cortical/amygdal/hippocampal/VTA glutamatergic projection neurons are dysregulated, creating DEGs at the synapse in the NAc? This part of the discussion should be more specific on what they mean.

We thank the reviewer for this comment. We agree that the description of the NAc circuitry was not connected to our own findings, and we have substantially shortened it, retaining only a single sentence summarizing the glutamatergic inputs to the NAc and its outputs to downstream regions (lines 351–356).

We have also revised the concluding sentence of this paragraph so that it follows from the preceding discussion of our DEGs (lines 351–356). Since our data indicate that glutamatergic synapse-associated genes are downregulated in NAc neurons after jSI, and since the NAc integrates glutamatergic inputs from several regions implicated in social and emotional behavior, we now state that altered glutamatergic signaling at these synapses may contribute to the behavioral abnormalities induced by jSI, rather than asserting that such deficits mediate them.

#3-minor-discussion-3
"Deficiencies in these proteins are associated with various behavioral abnormalities (Araujo et al., 2017; Chasse et al., 2024; Guo et al., 2020; Huang et al., 2021; Mukai et al., 2019)." line 334-335: what proteins and what behavioral abnormalities? This is not precise enough and needs reformulation/conclusions.

We thank the reviewer for this comment. We agree that the original sentence was not sufficiently specific, since it referred to deficiencies in several proteins and to behavioral abnormalities without indicating which protein was associated with which phenotype.

We have revised this passage to describe the reported phenotypes individually for each factor (lines 362–370). We have also incorporated here the background material on Setd1a and Brd4 that we removed from the Results section, as described in our response to the Reviewer #3's comment (#3-minor-results-3), so that the association of these factors with neurological and psychiatric conditions is presented in the Discussion rather than interrupting the presentation of our findings.

#3-minor-discussion-4
"A previous report suggests that histone modifications such as H3K4me3 in the hippocampus respond to an enriched environment (Schaffner et al., 2023), and our results indicate that these histone modifications may influence gene expression in the NAc under jSI stress as well." line 342-344: In which direction is the modification in the hippocampus in enriched environment? Is it opposite to what the authors have found in jSI (which would be interesting, since one could see a more social environment as an enriched condition too)? The idea behind this sentence needs to be precised.

We thank the reviewer for this comment. We have revised this sentence to describe the reported finding more precisely (lines 376–379), namely that the loss of H3K4me1 observed in SNCA transgenic mice was partially dampened by environmental enrichment.

Regarding the reviewer's suggestion that enrichment might be viewed as the converse of isolation, we agree this is an interesting possibility, but we have chosen not to develop the comparison in the text. Social isolation and environmental enrichment are not straightforwardly opposite conditions, since the absence of social contact is not simply the inverse of enrichment relative to standard housing, and the two may engage distinct circuits and cell populations. We therefore refer to this study only as evidence that these histone modifications are responsive to the housing environment, rather than drawing a directional comparison with our own data.

#3-minor-discussion-6
"Since neural development relies on the regulation of gene expression (Jain et al., 2001; Xiang et al., 2020), we hypothesize that these terms reflect altered gene expression regulation mechanisms under jSI stress." "However, these hypotheses need to be further validated by additional experiments." line 369-371 & 375-377: The authors are not integrating their results, they are being cautious, but the message stays unclear to the reader. What is the message here?

We thank the reviewer for this comment. We agree that our original wording was cautious to the point of leaving the message unclear, and we have rewritten this passage.

We now state explicitly what we wish to propose: that genes involved in transcriptional and chromatin regulation carried altered histone modifications, that changes at such loci may have consequences extending beyond the genes themselves through their downstream targets, and that this may be particularly relevant during the developmental window examined here (lines 404–416). The paragraph now concludes by presenting this as a hypothesis, namely that histone modification changes at regulatory genes act as an upstream event after jSI whose consequences are amplified through the targets of those regulators, together with a statement that this remains to be tested experimentally, rather than ending with a general remark that further validation is required.

#3-minor-discussion-8
"To determine whether histone modifications regulate specific genes, we focused on potentially important genes. Grik1, for example, exhibits reduced H3K27ac levels. It encodes a subunit of ionotropic glutamate receptors, and its deficiency has been found in mental diseases, such as schizophrenia and ADHD (Chatterjee et al., 2022; Hirata et al., 2012). The inactivation of Grik1 in rodents promotes anxiety-like behaviors via glutamatergic transmission (Englund et al., 2021)." line 387-392: What is the conclusion/hypothesis on Grik1's role?

We thank the reviewer for this comment. We agree that the original passage described what is known about *Grik1* without stating what we ourselves wished to conclude.

We have added a statement of our hypothesis at the end of this passage: that the reduction in H3K27ac around the *Grik1* locus contributes to the downregulation of *Grik1* after jSI, which in turn may contribute to the anxiety-like phenotypes associated with isolation (lines 432–437). We also indicate what would be required to test this, namely manipulating H3K27ac at the *Grik1* locus and assessing the resulting transcriptional and behavioral changes.

#3-minor-discussion-9
"Bcl2, for example, has downregulated H3K4me1, H3K4me3, and upregulated H3K27me3 levels. Bcl2 is an apoptosis-related gene that determines neuronal survival under stress. A previous study suggests that chronic social defeat stress decreases the Bcl-2/Bax ratio in NeuN+ neurons in the hippocampus (Zhu et al., 2024). Our data suggest that jSI is another type of stress that suppresses Bcl2 expression, and that the epigenetic factors are possible upstream

regulatory mechanisms." line 393-399: No links or clear hypothesis have been made here. The authors proposed to go deeper in this direction later. It would be interesting to conclude on the hypothesis on these two genes (Grik1/Bcl2) in their model.

We thank the reviewer for this comment. We agree that this passage describes our observations concerning *Bcl2* without stating what we conclude from them.

We have revised it to present our hypothesis explicitly, namely that the coordinated reduction of H3K4me1 and H3K4me3 together with the increase in H3K27me3 around the *Bcl2* locus contributes to its downregulation after jSI, and that this may in turn affect neuronal survival under stress (lines 443–446). We have also indicated what would be required to test this, in the same way as for *Grik1*, as described in our response to the Reviewer #3's comment (#3-minor-discussion-8).

#3-minor-discussion-10

"We found that Kdm6b may be at least partially involved in the gene expression changes in jSI mice, especially for potentially important genes like Nr4a1. Nr4a1 encodes a transcription factor that regulates dopamine metabolism, and it is reported to be involved in drug addiction, which is probably mediated by histone modification alterations." line 408-411: The authors are very cautious about their conclusion, maybe too much. Since they base their hypothesis on P14 cerebellum neurons in culture, more work is needed here to establish clearly the role of Kdm6b. It feels like the authors are dropping clues for the reader, without concluding themselves on their hypothesis.

We thank the reviewer for this comment, which we found particularly helpful. We recognize that our repeated use of hedging language left the reader uncertain as to what we were actually proposing.

We have restructured these passages so that our hypothesis is stated explicitly, followed by a statement of what would be required to test it (lines 455–469). For Kdm6b, we now propose that Kdm6b-mediated H3K27 demethylation contributes to the upregulation of *Nr4a1* in the NAc after jSI, and that testing this hypothesis will require NAc-specific manipulation of Kdm6b together with assessment of both transcriptional and behavioral outcomes. The limitations of the published dataset from which this hypothesis derives, including its origin in cerebellar tissue at P14, are stated in the Limitations section, as described in our response to the Reviewer #3's comment (#3-major-5).

We believe this approach addresses the reviewer's concern without overstating our findings, and it is consistent with the request from Reviewer #2 that causal language be moderated (#2-major-1).

#3-minor-discussion-11

"Our experiments suggest that Dkk3 is regulated by jSI stress, and this is probably mediated by Brd4." / "Our analysis also implies that Dkk3 and Hcrtr2 are probably regulated by other epigenetic regulators such as Setd1a." line 429-430 & 431-432: "Probably". No proof is given on that statement. Everything is "potential" or "possible" or "probable" here.

We agree with the reviewer. As described in our response to the Reviewer #3's comment (#3-minor-discussion-10), we have gone through the Discussion and removed or replaced the repeated use of "probably", "possible", and "potential", restructuring the relevant passages so that each hypothesis is stated explicitly, together with a statement of what would be required to test it (lines 476–483, 484-496).

#3-minor-discussion-12
"To summarize, we revealed the changes in gene expression and histone modification levels under jSI stress." line 444: Here they are missing the term "in the NAc" and "in female mice" and maybe need to add "some changes" in the sentence.

We thank the reviewer for this comment. We have revised the summary sentence to specify both the brain region and the sex of the animals studied, and to indicate that we identified a subset of changes rather than a comprehensive account (lines 498–499). It now reads: "To summarize, we revealed some changes in gene expression and histone modification levels in the NAc of female mice under jSI stress."

#3-minor-discussion-13
Major issue of this study is stated as "Another limitation related to this is that we inferred candidate epigenetic factors based on previously published public data. However, different brain regions, cell types, and experimental conditions between our analyses and public datasets may contribute to the gene expression differences, leading to false negative or false positive results in this study." line 456-460: Indeed, as they highlight here, the differences between the datasets chosen and their initial context (jSI, NAc) is huge. Maybe the authors could explain better why did they choose these datasets instead of more similar ones.

Also, in the RNAseq / cut&tag analysis, they "were unable to separate these subtypes (D1+ or D2+) for this current analysis". This analysis could be interesting to see in the supplementary or a brief description of what they have tried in the discussion, since the composition of the NAc is made of about 90-95% of MSNs and a plethora of interneurons (Parvalbumin, Cholinergic, etc), leading to different circuit connectivity and function. The striatum also contains a third population of D1/D2 hybrid neurons, maybe this population (about 5-10% of the whole striatum) made the identification of the neuronal subtypes harder.

We thank the reviewer for these comments.

Regarding the choice of public datasets, we have revised the Limitations section to state that these datasets were selected as the closest available to our system, since no dataset in which these factors had been perturbed exists for the NAc or striatum, and we have added developmental stage and sex to the list of differences between those datasets and our own (lines 506–527).

Regarding the separation of neuronal subtypes, our nuclei were sorted using an anti-NeuN antibody, which labels neuronal nuclei broadly and does not distinguish D1- from D2-

positive medium spiny neurons. We did not attempt to separate these subtypes, and we have now stated this reason explicitly in the Limitations section. As the reviewer notes, this is a meaningful limitation, since changes restricted to one subtype, or occurring in opposite directions between subtypes, would be underestimated or missed entirely in our analysis. We agree that subtype-resolved approaches will be required to address this in future work.

#### 4. Description of analyses that authors prefer not to carry out

#2-major-4

Behavior is missing from this study. The Introduction says jSI affects "motor, emotional, learning, and sociability-related behaviors." This manuscript does not show that the molecular changes track those phenotypes in the same cohort. Without that link, the psychiatric disease framing stays speculative.

We thank the reviewer for this comment, and we apologize that our reasoning was not made clear in the original manuscript.

Behavioral phenotypes produced by this isolation paradigm have been characterized in our previous study (Sazhina et al., ***Neuroimage***, 2025), which used exactly the same protocol as the present work, with isolation from P21 to P35 and regrouping from P35 to P49. We have described these findings in the Introduction so that the behavioral consequences of our paradigm are explicit (lines 554–557), rather than referring only to the literature in general terms.

We did not perform behavioral testing on the animals used for molecular analysis, because behavioral testing itself constitutes a substantial stimulus. Exposure to a novel environment, handling, learning experience, and aversive stimuli such as the foot shock used in fear conditioning all induce transcriptional and epigenetic changes in the brain, including the induction of immediate early genes such as *Nr4a1* and *Fosl2*, which are among the candidate genes discussed in this study. Had the same animals been subjected to behavioral testing, we would not have been able to attribute the observed changes to jSI rather than to the testing procedure. We therefore used dedicated cohorts for molecular profiling, and we have stated this rationale explicitly in the revised Methods.

We acknowledge, nevertheless, that this design means we cannot demonstrate a correspondence between molecular changes and behavioral phenotypes within the same individuals. We have stated this limitation in the Limitation section and have moderated the framing of our findings in relation to psychiatric disease accordingly (lines 524–527), as described in our response to the Reviewer #2's comment (#2-major-1).

#3-major-3（OPTIONAL）

Males: Experiments are only focused on females here. The authors didn't state why. These experiments (RNAseq, Cut&Tag) could be performed independently in male mice. The introduction refers to several pathologies that show sex-bias in human or animal models and even articles with sex differences. If the authors had a reason to select female only, it should be clearly stated. This proposition would take the same amount of resources and time that for this issue but would increase the knowledge about the (sex differences in the) effect of juvenile social isolation greatly.

We thank the reviewer for this suggestion, and we agree that a parallel analysis in male mice would substantially extend the value of this work.

As described in our response to the Reviewer #1's comment (#1-6), our focus on female mice was not arbitrary. In our previous study using the same isolation paradigm (Sazhina et al., ***Neuroimage***, 2025), jSI produced a heightened fear response in female but not male mice, and the present study was designed to examine the molecular basis of that female-specific phenotype in the NAc. We apologize that this rationale was not stated in the original manuscript, and we have described it explicitly in the Introduction and Methods of the revised version (lines 73–74, 524-527, 542-543).

We would nevertheless like to explain why we are unable to perform the proposed experiments within the scope of this revision. Generating a comparable dataset in males would require a new cohort of animals, isolation and regrouping over four weeks, nuclear isolation and sorting, and the preparation and sequencing of five libraries per animal, followed by the full analysis pipeline. As the reviewer notes, this would require resources and time comparable to those invested in the present study, and it is not feasible within the revision period. We have stated in the Limitations section that a parallel analysis in males is required to distinguish shared mechanisms of jSI from sex-specific ones (lines 524–527), and we intend to pursue this in future work.

#3-major-4（OPTIONAL）

Behavior abnormalities: Showing behavior abnormalities after jSI of female mice or, if it has been published somewhere else previously, a general description of the phenotypes observed in juvenile and adult female mice. This experiment could be performed in one batch of female separated at P21 and regrouped at P35, with anxiety (openfield or elevated plus maze, 1 day each), sociability (direct or 3-Chambers, 1 day each) and eventually depressive-like behaviors/anhedonia (sucrose preference test, 1 week including habituation to the two bottles and/or tail suspension/forced swimming, 1 day) tested. Same tests would be performed in juveniles or in adults.

We thank the reviewer for this suggestion, and for noting that a description of previously published phenotypes would be an acceptable alternative to new behavioral experiments.

Behavioral phenotypes produced by this isolation paradigm have been characterized in our previous study (Sazhina et al., ***Neuroimage***, 2025), which used exactly the same protocol as the present work, with isolation from P21 to P35 and regrouping from P35 to P49. In that study, jSI produced a heightened fear response in female but not male mice. We have described these findings explicitly in the Introduction (lines 73–74), so that the behavioral consequences of our paradigm are stated rather than left to the reader to infer from the general literature.

As described in our response to the Reviewer #2's comment (#2-major-4), we deliberately did not perform behavioral testing on the animals used for molecular profiling, since behavioral testing itself induces transcriptional and epigenetic changes in the brain and would have confounded the changes attributable to isolation. We have stated this rationale explicitly in the revised Methods (lines 554–557).

#3-major-9（OPTIONAL）
Include more mechanistic experiments (as suggested in the discussion) on some of the factors identified (Kdm6b, Brd4, Setd1a) to better confirm their involvement in the jSI-induced changes in transcriptome (and behavior). Conditional KO in the NAc with viral infection (mouse line Kdm6b-flox or Brd4-flox or Setd1a-flox existing + AAV-cre) or AAV Crispr for specific knockdown could hardly be performed in the time window used in this study (at least 3week expression of the Cre/Cas9). But systemic or intracerebral pharmacological approach (ip injection of an inhibitor or cannula implantation, local intra-accumbens injection) could be performed in juvenile (1 week recovery post-surgery only, can be done at P21 before isolation).

We thank the reviewer for this thoughtful suggestion, and we agree that functional analysis of Kdm6b, Brd4, and Setd1a would substantially strengthen the conclusions of this study.

As the reviewer notes, conditional knockout approaches requiring viral expression are not feasible within the developmental window used here. Regarding the pharmacological alternatives proposed, systemic administration would not allow us to attribute any resulting changes specifically to the NAc, and local intra-accumbens administration, while addressing this point, would require establishing a new surgical and behavioral pipeline in juvenile animals alongside the molecular analyses. Neither is achievable within the revision period.

We have therefore stated explicitly in the Discussion that functional validation of these candidate factors is required to test the hypotheses raised here (lines 509–513), and we intend to pursue this in future work. We are grateful to the reviewer for these constructive suggestions, which we will take up in designing those experiments.

#3-minor-discussion-5
"Since our mice were isolated from P21 to P35, a period critical for the maturation of neurons (Makinodan et al., 2012; Walker et al., 2019; Yamaguchi et al., 2024), it is possible that the neuronal development process is affected by the isolated housing environment." line 351-354: This sentence states that juvenile social isolation during development could impair development. It is probable, since early life stress (even maternal stress) can have prolonged effect on the offspring behaviors. It is probable that the time-window of jSI is important for development.

OPTIONAL: The authors would benefit of more experiments here: 1) They could perform SI in the same conditions but in adult females and compare the DEGs observed in that case (less development related genes probably). 2) "Neurons during adolescence mainly experience synaptic pruning and elimination (Afroz et al., 2016; Germann et al., 2021; Watanabe & Kano, 2024), and isolation stress probably impairs such processes." Here, the author could check in the NAc of their jSI female mice the state of dendritic arborization of MSNs (number, length, etc) on NAc brain slices.

We thank the reviewer for these suggestions, both of which we agree would strengthen the study.

Regarding social isolation in adult females, a comparison with adult-isolated animals would indeed help to establish whether the changes we observed are specific to the adolescent period. This would require a new cohort, a full isolation and regrouping schedule, nuclear isolation and sorting, and library preparation and sequencing, and is not feasible within the revision period. We will note this as a direction for future work. We have, however, revised the Introduction to cite studies defining P21 to P35 as a critical period for maturation in this system (lines 64–65), so that the rationale for focusing on this window is better supported.

Regarding dendritic morphology, we agree that examining the arborization of NAc MSNs would provide a useful structural correlate of the developmental processes discussed in this section. We did not collect tissue suitable for morphological analysis from these animals, since the entire NAc punch was used for nuclear isolation, and this analysis would therefore also require a new cohort. We will likewise note this as a direction for future work, and we have revised the corresponding statements in the Discussion so that they are presented as hypotheses rather than as established consequences of isolation (lines 389–391).
